# The Integrator complex cleaves nascent mRNAs to attenuate transcription

**DOI:** 10.1101/748319

**Authors:** Deirdre C. Tatomer, Nathan D. Elrod, Dongming Liang, Mei-Sheng Xiao, Jeffrey Z. Jiang, Michael Jonathan, Kai-Lieh Huang, Eric J. Wagner, Sara Cherry, Jeremy E. Wilusz

## Abstract

Cellular homeostasis requires transcriptional outputs to be coordinated, and many events post transcription initiation can dictate the levels and functions of mature transcripts. To systematically identify regulators of inducible gene expression, we performed high-throughput RNAi screening of the *Drosophila* Metallothionein A (MtnA) promoter. This revealed that the Integrator complex, which has a well-established role in 3’ end processing of small nuclear RNAs (snRNAs), attenuates MtnA transcription during copper stress. Integrator complex subunit 11 (IntS11) endonucleolytically cleaves MtnA transcripts, resulting in premature transcription termination and degradation of the nascent RNAs by the RNA exosome, a complex also identified in the screen. Using RNA-seq, we then identified >400 additional *Drosophila* protein-coding genes whose expression increases upon Integrator depletion. We focused on a subset of these genes and confirmed that Integrator is bound to their 5’ ends and negatively regulates their transcription via IntS11 endonuclease activity. Many non-catalytic Integrator subunits, which are largely dispensable for snRNA processing, also have regulatory roles at these protein-coding genes, possibly by controlling Integrator recruitment or RNA polymerase II dynamics. Altogether, our results suggest that attenuation via Integrator cleavage limits production of many full-length mRNAs, allowing precise control of transcription outputs.

## INTRODUCTION

In response to physiological cues, environmental stress, or exposure to pathogens, specific transcriptional programs are induced (reviewed in Vihervaara et al. 2018). These responses are often coordinated, rapid, and robust, in part because many metazoan genes are maintained in a poised state with RNA polymerase II (RNAPII) engaged prior to induction (reviewed in Mayer et al. 2017; Core and Adelman 2019). In addition to promoter-proximal pausing, there are many regulatory steps post transcription initiation that dictate the characteristics and fate of mature transcripts. For example, alternative splicing and/or 3’ end processing events can lead to the production of multiple isoforms from a single locus, and these transcripts can have distinct stabilities, translation potential, or subcellular localization (reviewed in Braunschweig et al. 2013; Tian and Manley 2017).

It is particularly important that genes produce full length, functional mRNAs, and mechanisms such as telescripting, involving U1 snRNP, actively suppress premature cleavage and polyadenylation events in eukaryotic cells (Kaida et al. 2010; Berg et al. 2012; Venters et al. 2019). Nevertheless, many promoters are known to generate short unstable RNAs (Kapranov et al. 2007; Xu et al. 2009; Porrua and Libri 2015). This suggests that premature transcription termination may often occur, thereby limiting RNAPII elongation and production of full-length mRNAs (reviewed in Kamieniarz-Gdula and Proudfoot 2019). Moreover, this process can be regulated (Brannan et al. 2012; Wagschal et al. 2012; Chalamcharla et al. 2015; Chiu et al. 2018). For example, it was recently shown that the cleavage and polyadenylation factor PCF11 stimulates premature termination to attenuate the expression of many transcriptional regulators in human cells (Kamieniarz-Gdula et al. 2019). Potentially deleterious truncated transcripts generated by premature termination are often removed from cells by RNA surveillance mechanisms, including by the RNA exosome (reviewed in Zinder and Lima 2017; Kamieniarz-Gdula and Proudfoot 2019). However, the full repertoire of cellular factors and co-factors that control the metabolic fate of nascent RNAs, especially during the early stages of transcription elongation, is still unknown.

We thus performed an unbiased genome-scale RNAi screen in *Drosophila* cells to reveal factors that control the output of a model inducible eukaryotic promoter. Transcription of *Drosophila* Metallothionein A (MtnA), which encodes a metal chelator, is rapidly induced when the intracellular concentration of heavy metals (e.g. copper or cadmium) is increased (reviewed in Gunther et al. 2012) **(Fig. 1A)**. This increase in transcriptional output is dependent on the MTF-1 transcription factor, which re-localizes to the nucleus upon metal stress and binds to the MtnA promoter (Smirnova et al. 2000). Our RNAi screen identified MTF-1 and other known regulators of MtnA transcription (Marr et al. 2006), but also surprisingly identified the Integrator complex as a potent inhibitor of MtnA during copper stress. Integrator harbors an endonuclease that cleaves snRNAs and enhancer RNAs (Baillat et al. 2005; Lai et al. 2015), and we find that Integrator can likewise cleave nascent MtnA transcripts to limit mRNA production. Using RNA-seq, we find hundreds of additional *Drosophila* protein-coding genes whose expression increases upon Integrator depletion. Focused studies on a subset of these genes confirmed that Integrator can cleave these nascent RNAs, thereby limiting productive transcription elongation. Altogether, we propose that Integrator-catalyzed premature termination can function as a widespread and potent mechanism to attenuate expression of protein-coding genes.

**Figure 1.**
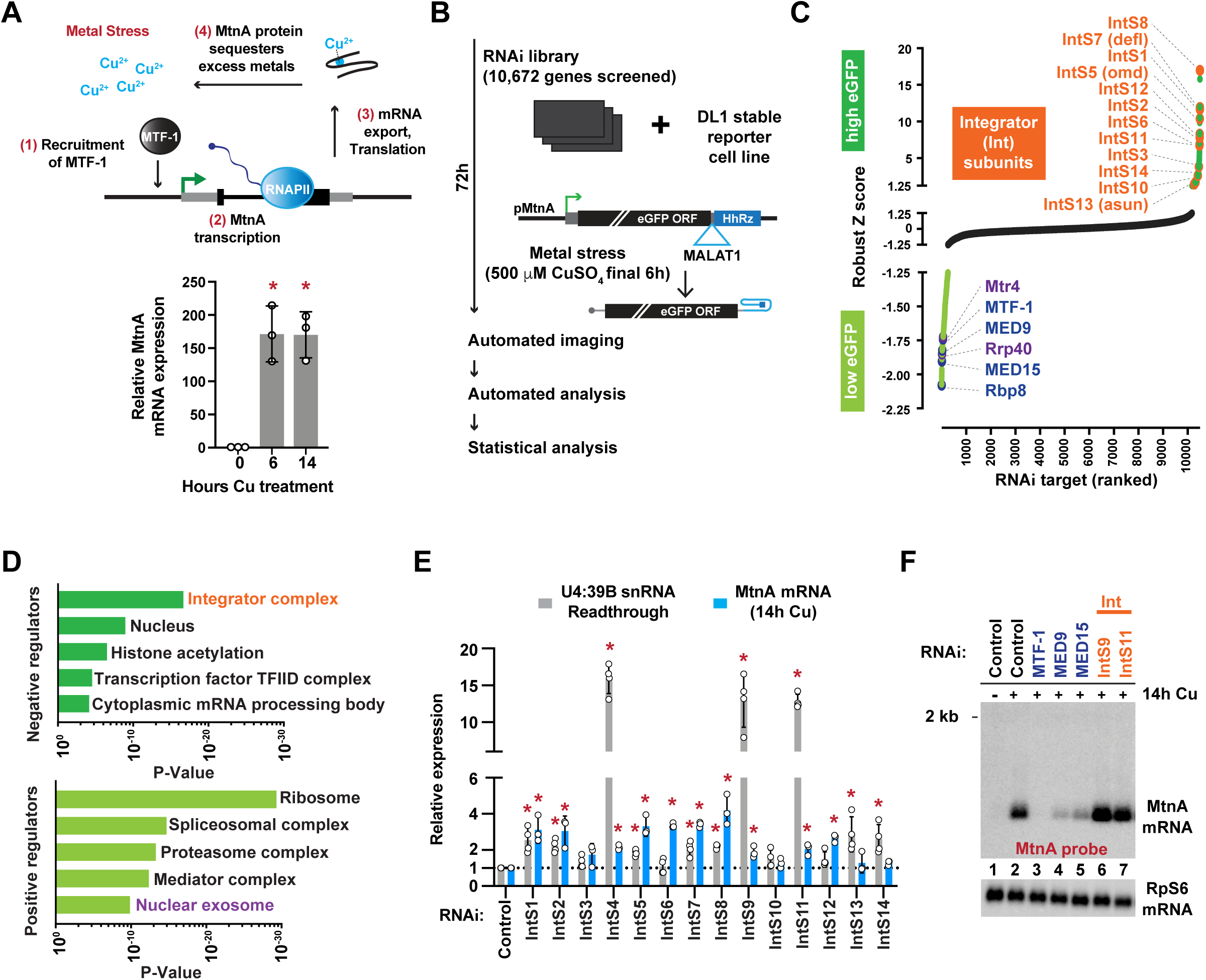
The Integrator complex inhibits expression from the MtnA promoter during copper stress. **(A)** *Top,* Upon metal stress, the transcription factor MTF-1 binds and induces transcription from the MtnA promoter, resulting in production of a protein that sequesters the excess metals to alleviate the stress. *Bottom, Drosophila* DL1 cells were treated with 500 µM copper sulfate (CuSO_4_) for the indicated times and RT-qPCR was used to measure endogenous MtnA mRNA expression. Data from 3 independent experiments were normalized to RpL32 mRNA expression and are shown as mean ± SD, *p<0.05. **(B)** RNAi screen pipeline using DL1 cells stably maintaining an eGFP reporter driven by the MtnA promoter. The self-cleaving hammerhead ribozyme (HhRz) (Dower et al. 2004) generates the eGFP mRNA 3’ end, which is then stabilized by the MALAT1 triple helix structure (Wilusz et al. 2012). **(C)** Robust Z scores of eGFP integrated intensity are shown. RNAi treatments that resulted in increased (Z score>1.3, dark green) or decreased (Z score<-1.3, light green) eGFP expression are marked, including Integrator subunits (orange), transcription regulators (blue), and RNA exosome components (purple). **(D)** Gene ontology (GO) analysis was performed to identify categories of genes that are enriched among the negative (Z score>1.3) and positive (Z score<-1.3) regulators of the eGFP reporter. **(E)** DL1 cells were treated with dsRNAs for 3 days to induce RNAi and depletion of the indicated factors. Expression of endogenous MtnA mRNA (after 14 h CuSO_4_ treatment) was quantified by RT-qPCR, and readthrough transcription downstream of the U4:39B snRNA was quantified by Northern blotting. Data are shown as mean ± SD, N≥3. *p<0.05. **(F)** Representative Northern blot of endogenous MtnA mRNA isolated from DL1 cells treated with dsRNA to induce RNAi of the indicated factor.

## RESULTS

### Genome-scale RNAi screening reveals the Integrator complex as a potent inhibitor of the MtnA promoter during copper stress

To identify regulators of an archetype inducible transcription program, the *Drosophila* MtnA promoter was cloned upstream of an intronless, non-polyadenylated eGFP reporter **(Fig. 1B)**. Maturation of this mRNA is independent of many canonical mRNA processing events, and we thus reasoned that high-throughput RNAi screening using this reporter should primarily identify transcriptional and translational regulators. *Drosophila* DL1 cells stably maintaining the reporter were treated with double-stranded RNAs (dsRNAs) for 3 days and copper was added for the final 6 h to activate the MtnA promoter and eGFP expression **(Fig. 1B)**. Automated microscopy and image analysis was then used to quantify eGFP fluorescence. 232 factors were required for eGFP expression during copper stress, including ribosomal subunits and well-characterized transcriptional regulators such as RNAPII, Mediator subunits (e.g. MED9 and MED15), and the MTF-1 transcription factor (Marr et al. 2006; Gunther et al. 2012) **(Fig. 1C, D; Supplemental Table S1)**. Unexpectedly, nearly all 14 subunits of the Integrator (Int) complex (reviewed in Baillat and Wagner 2015) were among the most potent negative regulators identified **(Fig. 1C, D)**. Of the >10,000 genes screened, depletion of the IntS8 subunit resulted in the largest increase in eGFP expression.

RT-qPCR and Western blotting confirmed that the dsRNAs resulted in depletion of the Integrator subunits **(Supplemental Fig. S1)** and increases in eGFP reporter expression at both the protein **(Supplemental Fig. S2A-C)** and mRNA levels **(Supplemental Fig. S2D-E)**. These increases in expression upon Integrator depletion were observed regardless of the ORF downstream of the MtnA promoter, as we also observed an increase when eGFP was replaced with nano-Luciferase **(Supplemental Fig. S2F)**. Likewise, the increases in expression were not dependent on the mRNA 3’ end processing signals downstream of the ORF, as similar increases were observed when a canonical polyadenylation signal was present **(Supplemental Fig. S2A-C)**.

### The Integrator complex is present at the endogenous MtnA locus during copper stress and represses MtnA pre-mRNA levels

The Integrator complex has been implicated in a myriad of diseases, interacts with RNAPII, and contains the IntS11 RNA endonuclease that generates the 3’ ends of spliceosomal snRNAs (Baillat et al. 2005; Baillat and Wagner 2015). We confirmed that depleting subunits of the Integrator cleavage module (IntS4, IntS9, or IntS11) resulted in increased snRNA readthrough transcription (Ezzeddine et al. 2011; Albrecht et al. 2018) **(Fig. 1E; Supplemental Fig. S3A)**. Nevertheless, because mature snRNAs have long half-lives (Fury and Zieve 1996), their levels only marginally decreased over the 72 h timecourse of the experiment **(Supplemental Fig. S3B)**.

Integrator has also been implicated in transcription regulation at enhancers and at a subset of EGF-responsive mRNAs (Gardini et al. 2014; Lai et al. 2015), so we hypothesized that the Integrator complex could be directly acting at the MtnA promoter. Indeed, depletion of Integrator subunits resulted in increased expression of the endogenous MtnA mRNA **(Fig. 1E, F; Supplemental Fig. S3C)** and pre-mRNA **(Fig. 2A)** to a similar extent during copper stress. Transcription of the MtnA locus is also induced by cadmium stress **(Fig. 2B)**, but we found that depletion of Integrator subunits strikingly had no effect on MtnA pre-mRNA levels under these conditions **(Fig. 2C)**. This indicates that the Integrator complex can regulate the output of a protein-coding gene in a context-specific manner.

**Figure 2.**
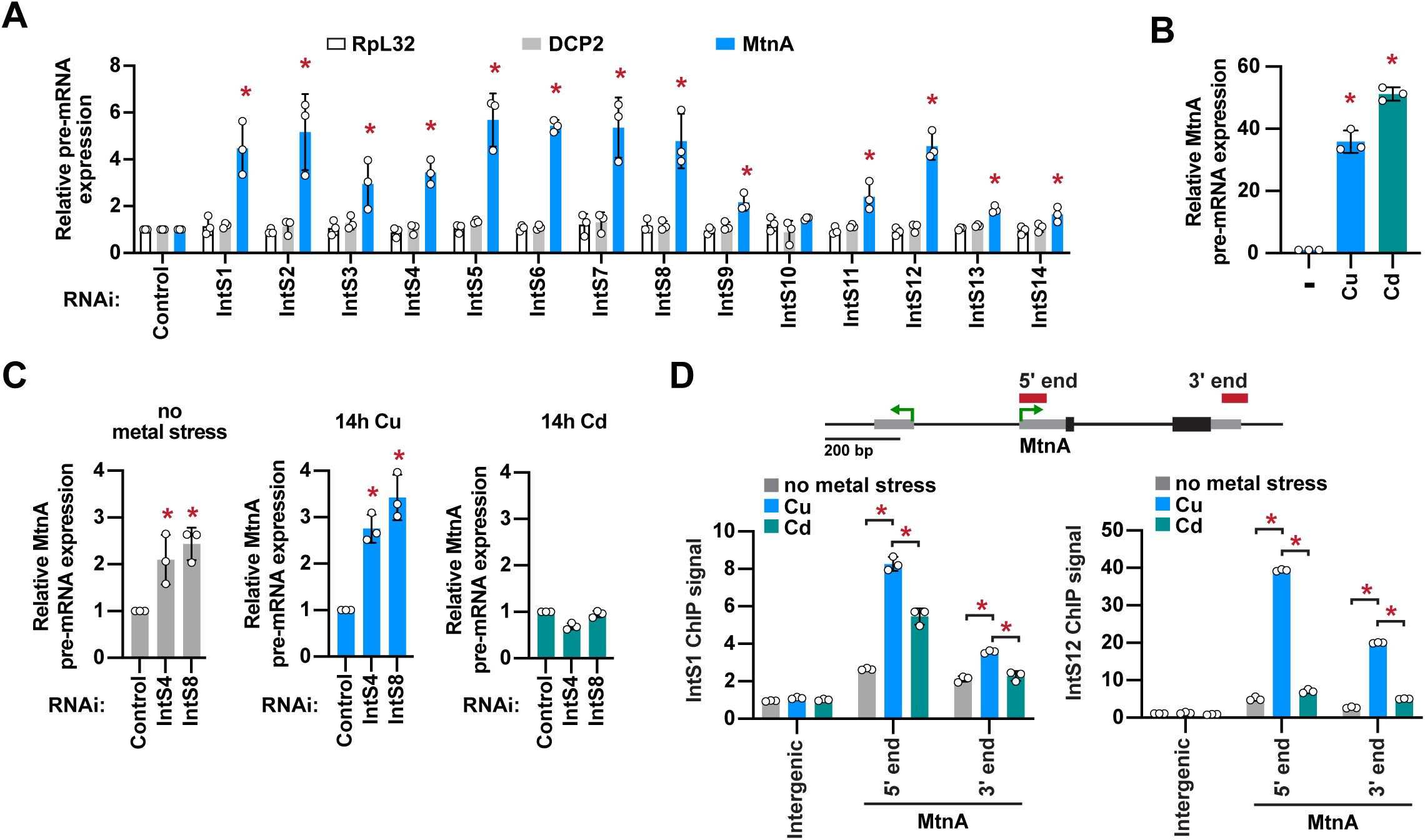
The Integrator complex is present at the MtnA locus during copper stress and attenuates MtnA transcription. **(A)** DL1 cells were treated with dsRNAs for 3 days to induce RNAi and depletion of the indicated factors. 500 µM CuSO_4_ was added for the last 14 h. RT-qPCR was then used to measure the pre-mRNA levels of endogenous RpL32, DCP2, and MtnA. Primer pairs that span an intron-exon boundary were used to specifically amplify pre-mRNAs. Data were normalized to RpL32 mRNA expression and are shown as mean ± SD, N=3. * p<0.05. **(B)** DL1 cells were unstressed (-) or treated with 500 µM CuSO_4_ or 50 µM CdCl_2_ for 14 h and RT-qPCR was then used to measure MtnA pre-mRNA levels. Data were normalized to RpL32 mRNA expression and are shown as mean ± SD, N=3. * p<0.05. **(C)** DL1 cells were treated with dsRNAs for 3 days and 500 µM CuSO_4_ or 50 µM CdCl_2_ was added for the final 14 h, as indicated. RT-qPCR was then used to measure MtnA pre-mRNA levels. Data were normalized to RpL32 mRNA expression and are shown as mean ± SD, N=3. * p<0.05. **(D)** The MtnA locus with the locations of ChIP amplicons. Recruitment of IntS1 and IntS12 in unstressed cells (gray) or after the cells had been treated with CuSO_4_ (blue) or CdCl_2_ (green) for 14 h was measured using ChIP-qPCR. Data are shown as fold change relative to the IgG control (mean ± SD, N=3). *p<0.05.

To explore the underlying basis for this distinct regulation of MtnA expression, we used chromatin immunoprecipitation (ChIP)-qPCR to examine the recruitment of Integrator subunits to the MtnA locus upon copper or cadmium stress. We found that IntS1 and IntS12 were recruited to the endogenous MtnA locus upon copper stress (especially to the 5’ end), but their recruitment was significantly less robust during cadmium stress **(Fig. 2D)**. These results suggest that Integrator-mediated repression of MtnA during copper stress is due to Integrator recruitment to the chromatin eliciting direct effects on transcription.

### The IntS11 endonuclease activity is required for Integrator-dependent regulation of MtnA expression

To define how Integrator functions at MtnA during copper stress, we first addressed whether the RNA endonuclease activity of IntS11 is required. The endogenous IntS11 protein was depleted using a dsRNA targeting either the IntS11 ORF or 3’ untranslated region (UTR) **(Fig. 3A)**, and this resulted in increased expression of endogenous MtnA mRNA relative to treatment with a control dsRNA **(Fig. 3B, lanes 1-3)**. Expression of a wildtype (WT) IntS11 transgene **(Fig. 3A)** in cells treated with the IntS11 3’ UTR dsRNA restored MtnA expression to levels similar to control treated cells **(Fig. 3B, lane 4 vs. 6)**, whereas expression of a catalytically dead (E203Q) (Baillat et al. 2005) IntS11 transgene did not **(Fig. 3B, lane 7 vs. 9)**. As a control, we confirmed that increased MtnA expression was observed when cells were treated with a dsRNA targeting the IntS11 ORF **(Fig. 3B, lanes 5 & 8)**, which depletes both endogenous and exogenously expressed IntS11 **(Fig. 3A)**. This requirement for IntS11 endonuclease activity was also observed with the eGFP reporter driven by the MtnA promoter **(Supplemental Fig. S4)**. Given that the E203Q mutation does not disrupt Integrator complex integrity (Baillat et al. 2005), these collective results indicate that the IntS11 endonuclease activity is required for Integrator to negatively regulate the transcriptional output of the MtnA promoter.

**Figure 3.**
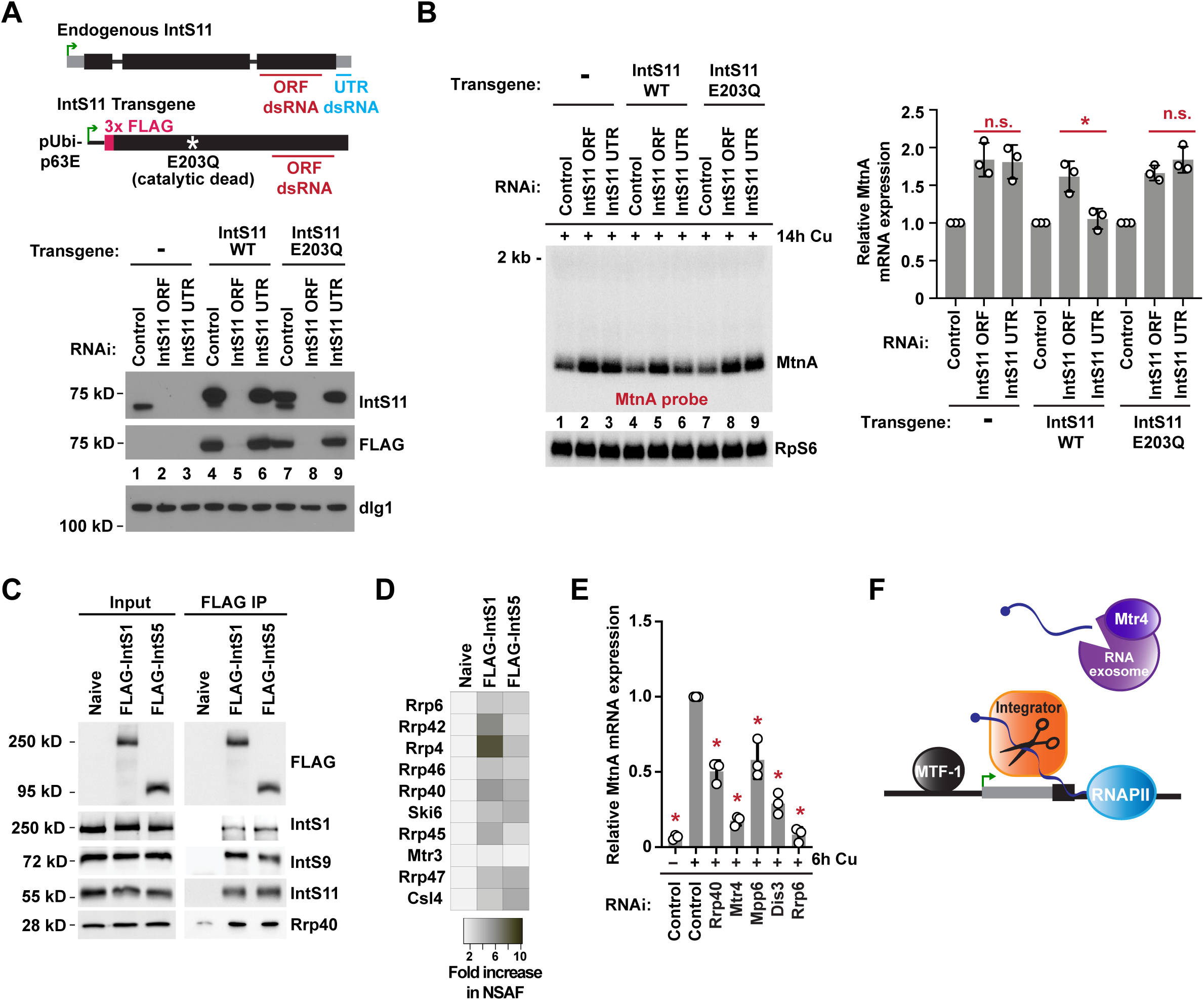
The IntS11 endonuclease activity and the RNA exosome regulate MtnA transcript levels. **(A)** *Top,* Schematic of IntS11 knockdown/plasmid rescue strategy. The ORF dsRNA (red) depletes IntS11 generated from both the endogenous locus and transgenes that are driven by the ubiquitin-63E promoter (pUbi-p63E), while the UTR dsRNA (blue) only depletes endogenous IntS11. *Bottom,* DL1 cells (lanes 1-3) or DL1 cells stably maintaining FLAG-tagged wildtype (WT; Lanes 4-6) or catalytically dead (E203Q; Lanes 7-9) IntS11 transgenes were treated with the indicated dsRNAs for 3 days. Western blot analysis was then used to examine IntS11 protein levels. dlg1 was used as a loading control. Representative blots are shown. **(B)** Representative Northern blot of the endogenous MtnA mRNA in DL1 cells stably maintaining IntS11 transgenes that had been treated with the indicated dsRNAs and CuSO_4_. Expression was quantified using ImageQuant and data are shown as mean ± SD, N=3. *p<0.05. **(C)** Western blot analysis of S2 nuclear extracts that were subjected to purification using FLAG affinity resin. Left panel, Western blots showing input levels of the denoted proteins in nuclear extracts derived from naïve S2 cells, S2 cells stably expressing FLAG-IntS1, or S2 cells stably expressing FLAG-IntS5. Right panel, same as left panel except that FLAG immunoprecipitates were probed. **(D)** Heat map showing the relative enrichment of RNA exosome components within FLAG-Integrator subunit purifications as determined by mass spectrometry. Normalized Spectral Abundance Factor (NSAF) values were quantified as previously described (Zybailov et al. 2006) and fold-enrichment of each RNA exosome subunit in Integrator purifications relative to purification from naïve extract is given. **(E)** DL1 cells were treated with dsRNAs for 3 days to induce RNAi and depletion of the indicated factors. CuSO_4_ was added for the last 6 h. Northern blots were then used to quantify expression of endogenous MtnA mRNA and data are shown as mean ± SD, N=3. *p<0.05. **(F)** Model for Integrator-dependent premature termination. After cleavage of the nascent MtnA transcript by IntS11, the small RNAs are degraded by the RNA exosome.

### The Integrator complex cleaves nascent MtnA mRNAs to trigger transcription termination

By immunoprecipitating Integrator subunits followed by immunoblotting **(Fig. 3C)** or mass spectrometry **(Fig. 3D)**, we found that Integrator interacts with the nuclear RNA exosome, which catalyzes 3’-5’ degradation of many RNAs (reviewed in Zinder and Lima 2017). Interestingly, our genome-scale RNAi screen identified many RNA exosome core components and co-factors (including Rrp40 and Mtr4) as *positive* regulators of the MtnA promoter during copper stress **(Fig. 1C, D)**. This suggests an interplay between the RNA exosome and Integrator complex and that the RNA exosome may also function in controlling MtnA transcriptional output. We confirmed that RNA exosome core components and co-factors were efficiently depleted by RNAi **(Supplemental Fig. S5A)**, and that this resulted in the expected ribosomal RNA processing defects **(Supplemental Fig. S5B)** as well as decreased output from the MtnA promoter during copper stress **(Fig. 3E; Supplemental Fig. S5C, D)**. We thus hypothesized that Integrator may cleave nascent MtnA transcripts to prematurely terminate RNAPII transcription (as Integrator is enriched near the 5’ end of the MtnA genomic locus during copper stress **(Fig. 2D)**) and that these cleaved transcripts are targeted for rapid degradation by the RNA exosome **(Fig. 3F)**.

To test this model, we first treated cells with dsRNAs to deplete the exosome-associated RNA helicase Mtr4 (Lubas et al. 2011). This resulted in a reduction in full-length endogenous MtnA mRNA expression **(Fig. 3E; Supplemental Fig. S5E)** and, concomitantly, a number of small RNAs were detected from the MtnA locus, including prominent transcripts with lengths of ∼85 and ∼110 nt **(RNAs marked in orange; Fig. 4A, Lane 4)**. These small RNAs were dependent on the MTF-1 transcription factor **(Fig. 4A, Lane 6)** and capped at their 5’ ends, as they could be degraded by a 5’ phosphate-dependent exonuclease only after treatment with Cap-Clip Acid Pyrophophatase which hydrolyzes cap structures to generate 5’ monophosphate groups **(Supplemental Fig. S6A)**. Northern blots using multiple probes further indicated that the small RNAs have the same TSS as MtnA mRNA **(Supplemental Fig. S6B)** and ligation-mediated 3’ RACE revealed that these small RNAs had detectable 3’ oligoadenylation, a mark known to facilitate RNA degradation by the RNA exosome (LaCava et al. 2005) **(Supplemental Fig. S6C, D; Supplemental Table S2)**.

**Figure 4.**
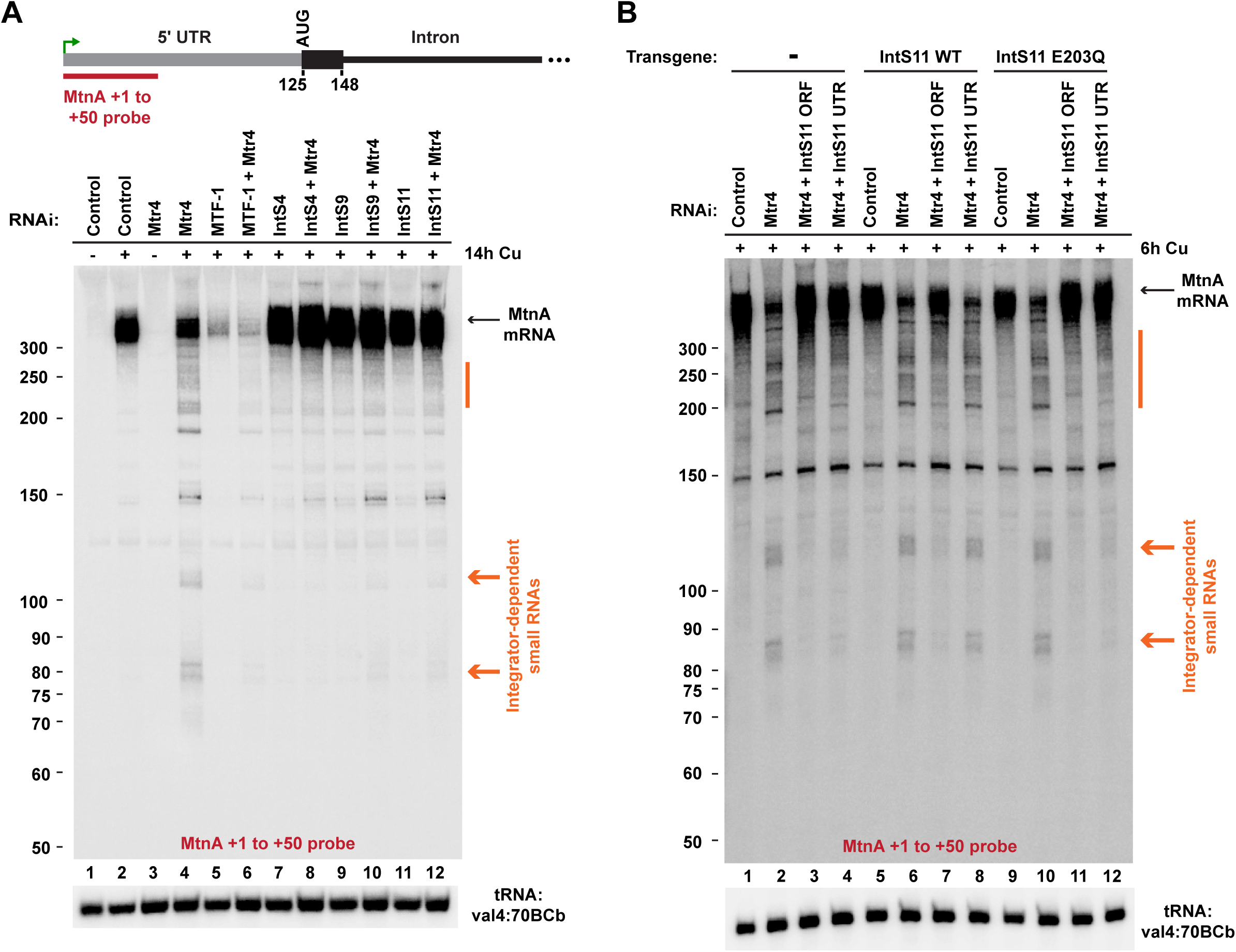
The Integrator complex cleaves nascent MtnA RNAs to catalyze premature transcription termination. **(A)** Northern blotting was used to analyze RNAs generated from the endogenous MtnA locus in DL1 cells treated with the indicated dsRNAs and CuSO_4_. Full-length MtnA mRNA (black arrow) and Integrator-dependent small RNAs (orange) are indicated. **(B)** Parental DL1 cells (Lanes 1-4) or DL1 cells stably expressing WT (Lanes 5-8) or catalytically inactive (Lanes 9-12) IntS11 transgenes were treated with the indicated dsRNAs and CuSO_4_. Northern blotting was then performed as in A.

We next co-depleted Mtr4 and Integrator subunits and observed that these small RNAs were completely eliminated and that full-length MtnA mRNA expression was restored **(Fig. 4A, Lanes 8, 10, & 12; Supplemental Fig. S5E)**. This suggests that generation of the small RNAs is dependent on Integrator. In support of this, in cells in which the endogenous IntS11 protein had been depleted by RNAi **(Fig. 4B, lane 4)**, re-expression of a wildtype (WT) IntS11 transgene restored expression of the MtnA small RNAs **(Fig. 4B, lane 8)**, whereas expression of a catalytically dead (E203Q) (Baillat et al. 2005) IntS11 transgene did not **(Fig. 4B, lane 12; Supplemental Fig. S7)**.

Together, the data in **Figs. 1-4** support a model in which the Integrator complex is recruited to the active MtnA locus during copper stress to cleave nascent RNAs, thereby facilitating premature termination **(Fig. 3F)**. This role for Integrator at MtnA is mechanistically related to its function at snRNA genes, where Integrator both cleaves the nascent snRNA and promotes RNAPII termination/recycling. Rather than Integrator attaining a novel function at protein-coding genes (Gardini et al. 2014; Stadelmayer et al. 2014; Skaar et al. 2015), our data show that the Integrator endonuclease activity has been ‘repurposed’ at the MtnA gene. Once generated, the prematurely terminated small RNAs are actively targeted for 3’-5’ degradation by the nuclear RNA exosome, at least in part due to 3’ oligoadenylation. Degradation of these small RNAs appears to be critical for enabling subsequent rounds of MtnA transcription, as the output of the MtnA promoter is reduced when the RNA exosome is depleted from cells **(Fig. 1C, 3E)**. The requirement of the RNA exosome for MtnA transcription is, however, abrogated when Intergator is depleted or catalytically inactive. These results indicate a potential epistastic relationship in which Integrator cleaves nascent RNAs that must be subsequently degraded to allow production of more full-length mRNA transcripts in copper stressed cells.

### Many *Drosophila* protein-coding genes are controlled by the Integrator complex

As Integrator-dependent termination events potently attenuate transcription from the MtnA promoter during copper stress, we next asked whether additional protein-coding genes are similarly regulated. IntS9, which forms a heterodimer with IntS11 and is essential for its endonuclease activity (Wu et al. 2017), was depleted from DL1 cells for 3 days and copper was added for the final 14 h. Using RNA-seq, we identified 409 and 49 genes that were up- and down-regulated, respectively, upon IntS9 depletion (fold change >1.5 and p<0.001) **(Fig. 5A; Supplemental Table S3)**. The set of up-regulated mRNAs was enriched in genes that respond to stimuli as well as gene ontology categories related to cell migration, proliferation, and cell fate specification **(Supplemental Fig. S8)**. In contrast, no gene ontology categories were enriched in the set of down-regulated mRNAs.

**Figure 5.**
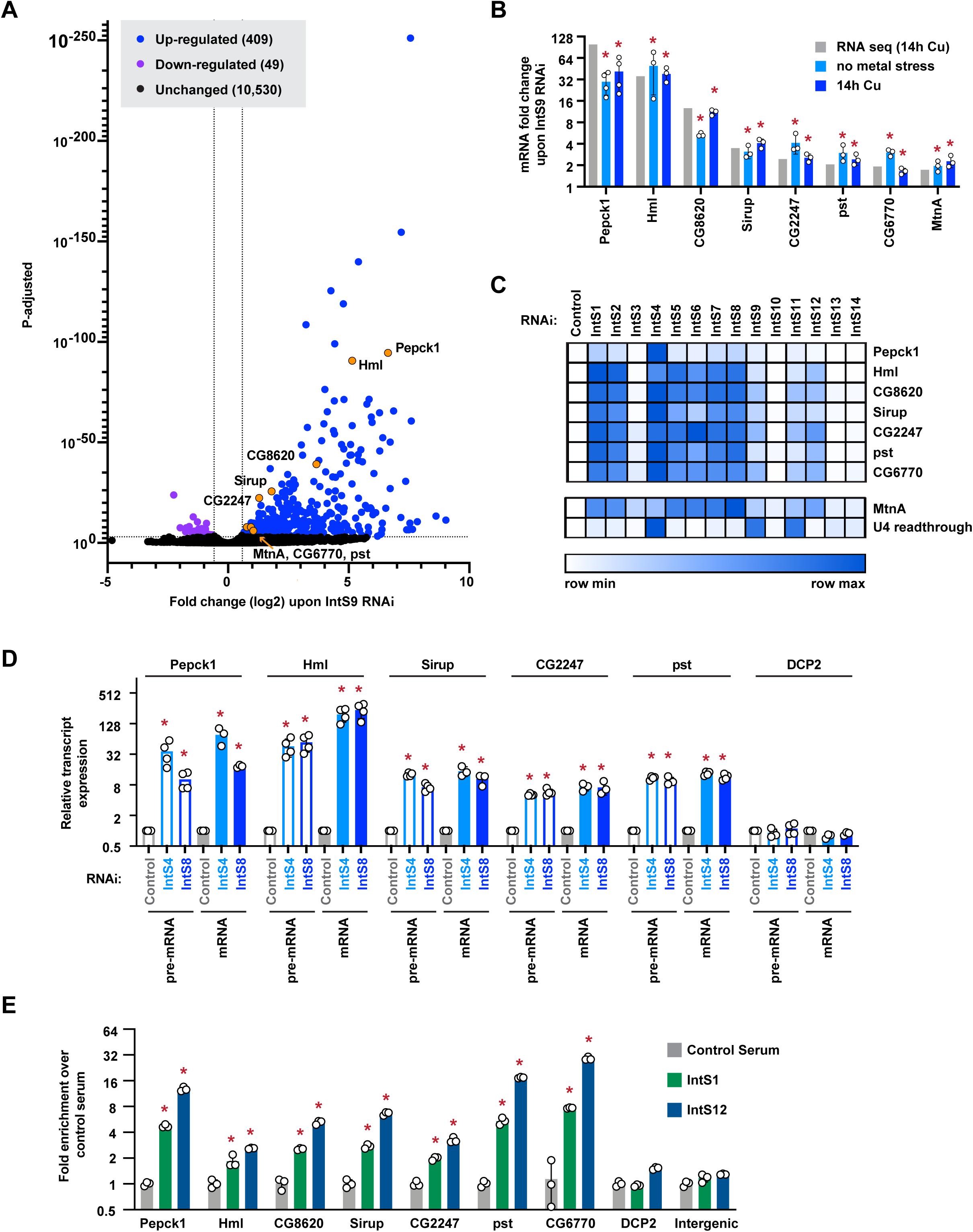
Integrator depletion results in up-regulation of many protein-coding genes. **(A)** DL1 cells were treated for 3 days with a control (βgal) dsRNA or a dsRNA to deplete IntS9 and CuSO_4_ was added for the last 14 h. Total RNA was isolated, depleted of ribosomal RNAs, and RNA-seq libraries prepared (3 biological replicates per condition). The mangnitude of change in mRNA expression compared with statistical significance (P value) is shown as a volcano plot. Threshold used to define IntS9-affected mRNAs was fold change >1.5 and p<0.001. **(B)** To verify the RNA-seq results (gray), DL1 cells were treated for 3 days with a control (βgal) dsRNA or a dsRNA to deplete IntS9 with or without CuSO_4_ added for the last 14 h (light blue and dark blue, respectively). RT-qPCR was then used to quantify changes in mRNA expression levels. Data were normalized to RpL32 mRNA expression and are shown as mean ± SD compared to treatment with a control dsRNA, N≥3. *p<0.05. **(C)** DL1 cells were treated with dsRNA to induce RNAi of the indicated factor and RT-qPCR was then used to quantify changes in mRNA expression levels. CuSO_4_ was added for the last 14 h only when measuring MtnA mRNA levels. Northern blotting was used to quantify readthrough transcription downstream of the U4:39B snRNA as described in Fig. 1E. Data are summarized as a heat map using Morpheus (Broad Institute) with darker shades representing increased transcript expression compared to treatment with a control (βgal) dsRNA. Individual RT-qPCR data points are provided in Supplemental Fig. S9. **(D)** RT-qPCR was used to measure the mRNA and pre-mRNA levels of the indicated transcripts. Data were normalized to RpL32 mRNA expression and are shown as mean ± SD, N≥3. * p<0.05. **(E)** ChIP-qPCR was used to measure IntS1 and IntS12 occupancy at the indicated promoter regions. Data are shown as fold change relative to the IgG control serum (mean ± SD, N=3). *p<0.05.

To validate the RNA-seq results, seven mRNAs that had differing magnitudes of fold change upon IntS9 depletion were selected for further analysis (genes marked in orange in **Fig. 5A**). Among these genes, five contain introns and two are intronless (CG8620 and CG6770). RT-qPCR confirmed that expression of all seven of these mRNAs increased upon IntS9 depletion regardless of whether the cells were subjected to metal stress **(Fig. 5B)**. Therefore, we did not induce metal stress in subsequent experiments. Upon depleting each Integrator subunit individually, we noted that expression of these mRNAs was often most affected by depletion of IntS4, which forms the scaffold of the Integrator cleavage module (Albrecht et al. 2018) **(Fig. 5C; Supplemental Fig. S9)**. Moreover, analogous to our prior results with MtnA **(Fig. 1E)**, depletion of many non-catalytic Integrator subunits (notably IntS1, IntS2, IntS5, IntS6, IntS7, and IntS8) also caused large increases in the expression of these mRNAs **(Fig. 5C; Supplemental Fig. S9)**.

### The Integrator complex cleaves many nascent mRNAs to trigger transcription termination

To determine if Integrator controls the outputs of these protein-coding genes transcriptionally or post-transcriptionally, we measured pre-mRNA levels from the intron-containing genes. Expression of these pre-mRNAs increased upon depletion of Integrator subunits, and the observed fold changes are similar to the increases in mature mRNA levels **(Fig. 5D)**. These results mirror our findings at MtnA **(Fig. 2A)** and strongly suggest that the observed increases in mature transcript levels are due to transcriptional control by Integrator, and not due to indirect effects of Integrator functioning in snRNA processing **(Supplemental Fig. S3A, B)**. Moreover, ChIP-qPCR confirmed that multiple Integrator subunits are recruited to the 5’ ends of these gene loci **(Fig. 5E)**. As an additional control, we monitored the DCP2 locus and observed minimal Integrator binding to the gene **(Fig. 5E)** as well as no change in DCP2 pre-mRNA or mRNA levels upon Integrator depletion **(Fig. 5D)**.

To further confirm that Integrator regulation of these genes was driven by their promoters, we cloned each of these regions (along with a portion of the 5’ UTRs) upstream of an eGFP reporter **(Fig. 6A)**. Indeed, the expression of eGFP mRNA driven from each of the examined promoters was sensitive to the levels of Integrator subunits, including those in the Integrator cleavage module (especially IntS4) as well as many of the non-catalytic subunits **(Fig. 6A; Supplemental Fig. S10)**. Similar results were obtained with the MtnA driven eGFP reporter **(Fig. 6A)**, whereas a reporter plasmid that monitors Integrator activity downstream of an snRNA (Chen et al. 2013) displayed a distinct sensitivity pattern **(Fig. 6B)**. For example, IntS6 depletion caused up-regulation of the output from all the Integrator regulated protein-coding gene promoters **(Fig. 6A)**, but had minimal effect on the U4:39B snRNA readthrough reporter **(Fig. 6B; Supplemental Fig. S10)**. As an additional control, we confirmed that expression of an eGFP reporter driven by the ubiquitin-63E (Ubi-p63e) promoter did not increase upon depletion of any of the Integrator subunits **(Fig. 6A; Supplemental Fig. S10)**. This is consistent with the RNA-seq results that showed endogenous Ubi-p63e mRNA levels do not change upon Integrator depletion **(Supplemental Table S3)**.

**Figure 6.**
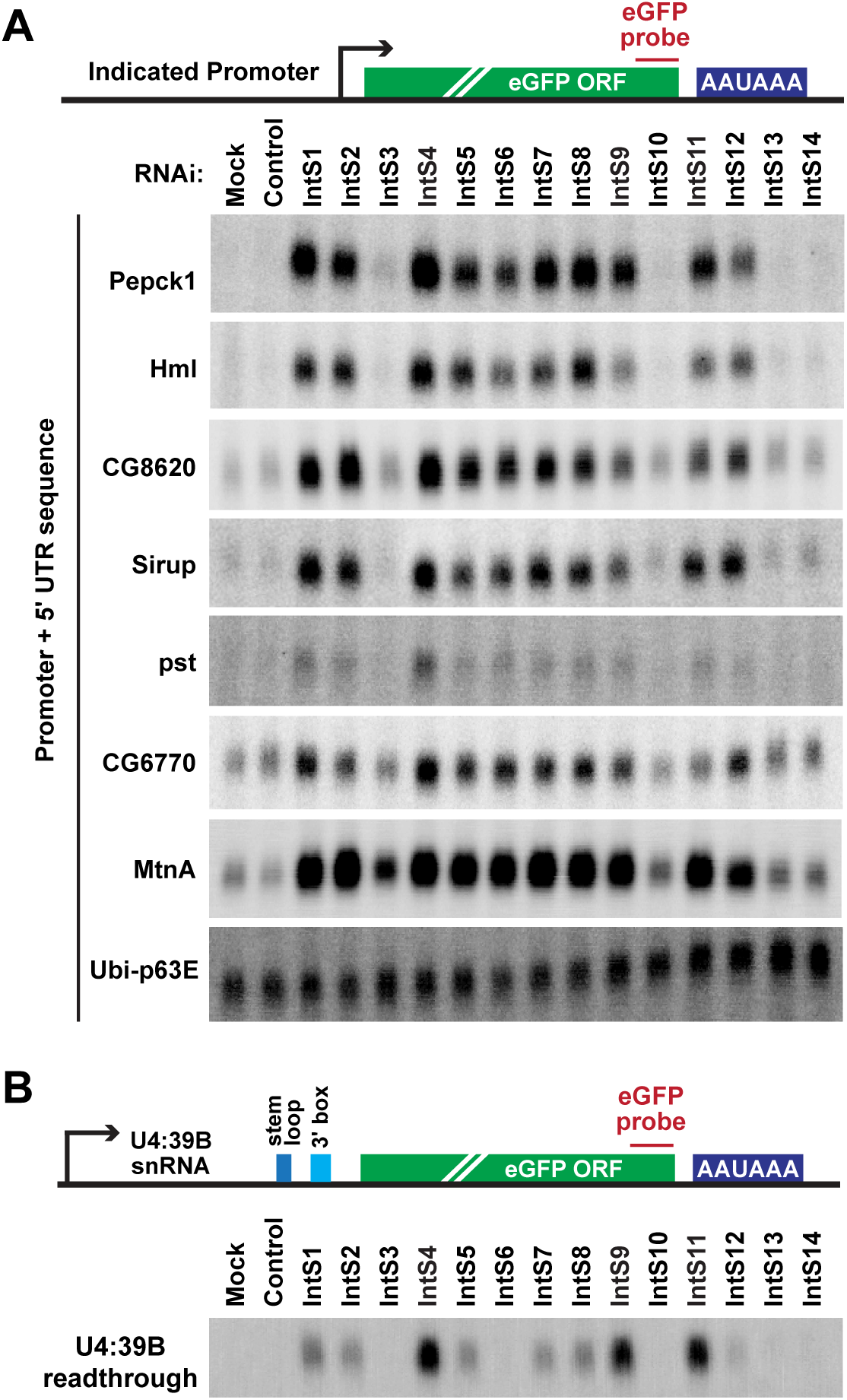
eGFP reporter genes driven by the exemplar promoters are regulated by Integrator. **(A)** The promoter and 5’ UTR of each of the indicated protein-coding genes was cloned upstream of an eGFP reporter. The plasmids were then individually transfected into DL1 cells that had been treated with the indicated dsRNAs. CuSO_4_ was added for the last 14 h only when measuring eGFP production from the MtnA promoter. Northern blots were used to quantify expression of each eGFP reporter mRNA. Representative blots are shown. Loading controls and quantification are provided in Supplemental Fig. S10. **(B)** DL1 cells were treated with the indicated dsRNAs and then transfected with a reporter plasmid that produces eGFP when the encoded U4:39B snRNA fails to be properly processed at its 3’ end. Northern blots were used to quantify eGFP mRNA expression that is a result of U4:39B readthrough. A representative blot is shown. Loading controls and quantification are provided in Supplemental Fig. S10.

Next, we tested whether Integrator catalyzes premature transcription termination at these genes in a manner analogous to how it controls MtnA. We first investigated if the IntS11 endonuclease activity is required. Depletion of the endogenous IntS11 protein using a dsRNA targeting the IntS11 3’ UTR **(Fig. 3A)** resulted in increased expression of each of the examined Integrator-dependent mRNAs **(Fig. 7A)**. Expression of a wildtype (WT) IntS11 transgene restored mRNA expression to levels similar to control treated cells, whereas the catalytically dead IntS11 E203Q mutant did not **(Fig. 7A)**. The IntS11 endonuclease activity is thus indeed required for regulation of each of these genes and, notably, the presence of the E203Q mutant protein exacerbated the changes in expression of the CG6770, pst, and Sirup mRNAs, potentially indicative of a dominant negative effect.

**Figure 7.**
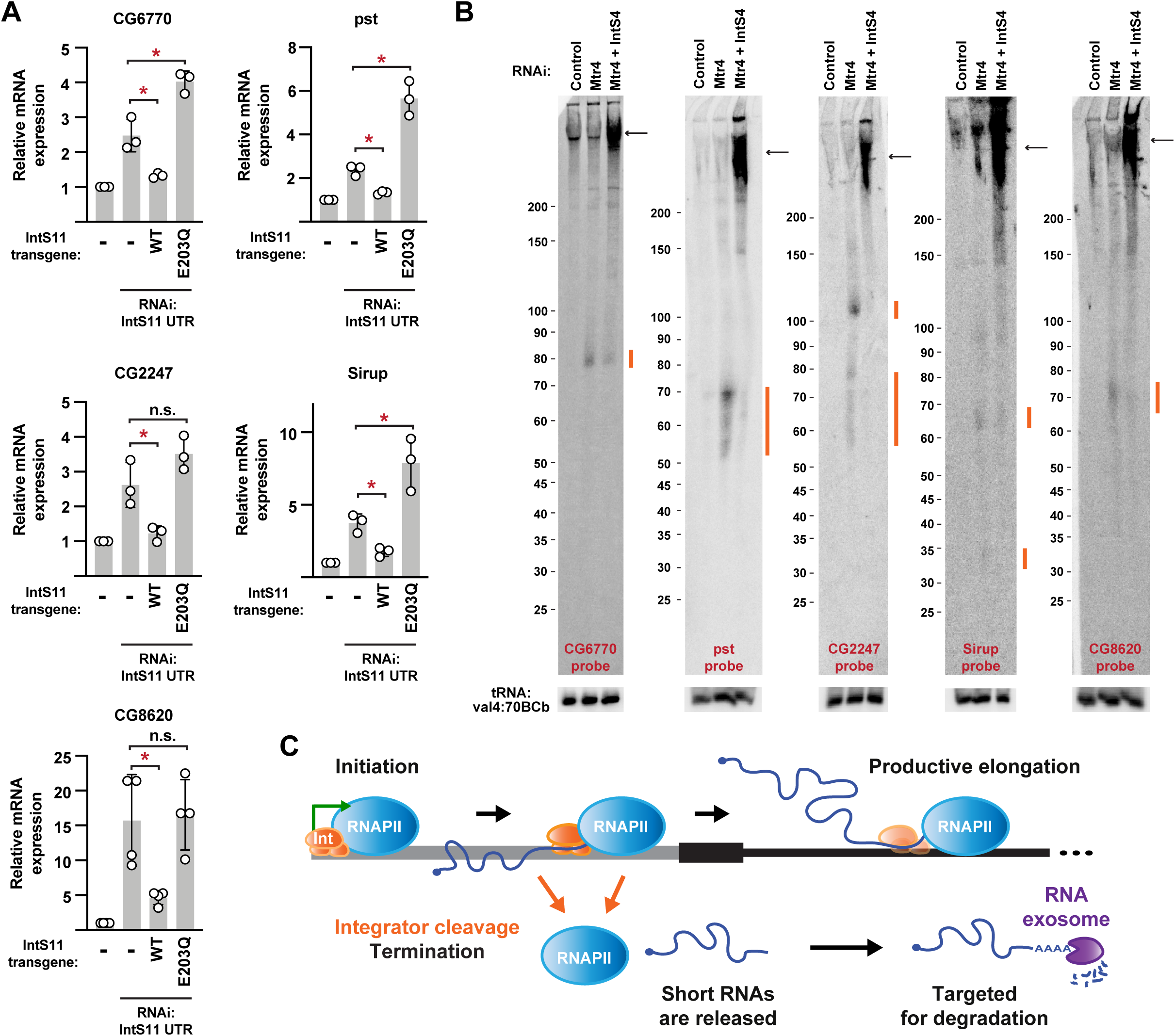
The Integrator complex cleaves many nascent mRNAs to catalyze premature transcription termination. **(A)** Parental DL1 cells (denoted -) or DL1 cells stably expressing WT or catalytically inactive (E203Q) IntS11 transgenes were treated with a dsRNA to the IntS11 3’ UTR, thereby depleting endogenous IntS11 but not IntS11 made from the transgenes. RT-qPCR was then used to quantify expression of the indicated mRNAs. Data were normalized to RpL32 mRNA expression and are shown as mean ± SD, N=3. * p<0.05. **(B)** DL1 cells were treated with dsRNAs for 3 days to induce RNAi and depletion of the indicated factors. Northern blotting using 50 μg of total RNA was then used to analyze transcripts from the 5’ ends of the indicated protein-coding loci. Integrator-dependent small RNAs (orange) and full-length mRNAs (black arrows) are noted. Representative blots are shown. **(C)** Schematic of a protein-coding locus, highlighting the presence of Integrator (Int, orange) and possible fates of RNAPII (blue). After transcription initiation, RNAPII can transition into productive elongation to generate the mature mRNA. Alternatively, Integrator can cleave the nascent RNA, thereby enabling transcription termination and degradation of the short RNA by the nuclear RNA exosome (purple).

Northern blots were then used to detect premature termination products from each of the Integrator regulated genes **(Fig. 7B)**. Given that the MtnA cleavage products are rapidly degraded by the RNA exosome **(Fig. 4)**, we reasoned that small RNAs generated from other loci would likewise be unstable. Depletion of the exosome-associated RNA helicase Mtr4 (Lubas et al. 2011) enabled small RNAs to be detected from the 5’ ends of the Integrator-dependent genes, and these transcripts were lost upon Integrator co-depletion **(RNAs marked in orange; Fig. 7B)**. Interestingly, these small RNAs were of defined lengths and often 50-110 nucleotides, roughly mirroring the sizes of cleavage products observed at the MtnA locus **(Fig. 4)**. We thus conclude that the Integrator complex is recruited to a number of protein-coding genes where it cleaves nascent RNAs and facilitates premature transcription termination **(Fig. 7C)**.

### Integrator cleavage of nascent mRNAs is likely independent of a 3’ box sequence

At snRNA gene loci, a conserved but relatively degenerate sequence known as the 3’ box is located 9-19 nt downstream of the 3’ ends of mature snRNA transcripts and is required for Integrator cleavage (Hernandez 1985; Baillat and Wagner 2015). A similar 3’ box-like sequence is not immediately recognizable within MtnA or the other transcripts we studied in detail, and we thus introduced deletions into the MtnA 5’ UTR upstream of the eGFP reporter in an attempt to alter the cleavage product sizes **(Supplemental Fig. S11A)**. Notably, this analysis revealed that Integrator-dependent small RNAs derived from the MtnA promoter appear to be largely 70-90 nt in length, regardless of the mRNA sequence **(Supplemental Fig. S11B)**. This suggests that Integrator may cleave nascent mRNAs at a set distance from the TSS in a manner independent of local DNA or RNA sequence content, perhaps at positions of RNAPII pausing/stalling or nucleosomes (Chiu et al. 2018).

## DISCUSSION

Altogether, our data indicate that the Integrator complex can attenuate the expression of protein-coding genes by catalyzing premature transcription termination **(Fig. 7C)**. The IntS11 endonuclease cleaves a subset of nascent mRNAs, which ultimately triggers degradation of the transcripts by the RNA exosome along with RNAPII termination. We suggest that many protein-coding genes are negatively regulated via this attenuation mechanism, and the *Drosophila* MtnA promoter highlights context-specific regulation by Intgerator. Transcription of MtnA is induced by copper or cadmium stress, and yet we find that Integrator is robustly recruited to the MtnA promoter only under copper stress conditions **(Fig. 2)**. This is not because the Integrator complex is generally diassembled or “poisoned” by cadmium, as Integrator continues to regulate the outputs of other protein-coding genes **(Supplemental Fig. S12)**. We instead propose that context-specific regulation of this locus may be related to the fact that cadmium is a strictly toxic metal, while copper is required for the function of a subset of enzymes and must be maintained in a narrow concentration range (reviewed in Festa and Thiele 2011). Therefore, homeostatic control of MtnA is required to maintain copper levels, while cells need to maximally produce MtnA in the presence of cadmium. We thus propose that regulation of MtnA levels by Integrator during copper stress is for fine-tuning purposes, perhaps to limit maximal transcriptional induction and/or facilitate transcriptional shut-off once copper stress has passed. Our results suggest that the Integrator complex can be recruited to gene loci only when needed, thereby ensuring tight control over transcriptional output.

In addition to cleaving MtnA transcripts, Integrator cleaves multiple other RNA classes in metazoan cells, including enhancer RNAs (Lai et al. 2015), snRNAs (Baillat et al. 2005), telomerase RNA (Rubtsova et al. 2019), and some herpesvirus microRNA precursors (Cazalla et al. 2011; Xie et al. 2015). Using RNA-seq, we expanded this list of Integrator target loci and identified hundreds of additional protein-coding genes that are negatively regulated by Integrator **(Fig. 5)**. We focused on a set of Integrator-dependent genes and found that Integrator catalyzes premature transcription termination of these genes **(Figs. 6, 7)**, which is consistent with prior studies that suggested roles for Integrator in termination (Skaar et al. 2015; Shah et al. 2018; Gómez-Orte et al. 2019). Some of these genes (CG8620, Pepck1, and Sirup) have promoter-proximal RNAPII that rapidly turns over (Shao and Zeitlinger 2017), which may indicate that Integrator can aid in clearing paused or stalled RNAPII. Once Integrator has cleaved the nascent mRNAs, we find that they are rapidly degraded from their 3’ ends by the RNA exosome **(Fig. 7C)**. This may be critical for enabling subsequent rounds of transcription (especially at the MtnA locus), perhaps because the small RNAs can form stable RNA-DNA hybrids (R-loops) that block transcription initiation or elongation (Belotserkovskii et al. 2017; Nojima et al. 2018; Crossley et al. 2019).

Endonucleolytic cleavage is critical for Integrator regulation at snRNA and protein-coding genes, but our data indicate that these loci have different dependencies on Integrator subunits. Our genetic studies indicate that Integrator subunits 4, 9, and 11 (which form the Integrator cleavage module) are most important for snRNA processing, while the non-catalytic Integrator subunits (all of which currently lack annotated molecular functions) play minor roles **(Fig. 1E, 6B)**. In contrast, large increases in mRNA expression were observed when many of the non-catalalytic subunits were depleted (especially IntS1, IntS2, IntS5, IntS6, IntS7, and IntS8) **(Fig. 1E, 5C, 6A)**. IntS13 was recently shown to be able to function independently from other Integrator subunits at enhancers (Barbieri et al. 2018), suggesting the existence of submodules or “specialized” complexes that may enable the activity and function of Integrator to be distinctly regulated depending on the gene locus and cellular state. Future work will reveal the subunit requirements of Integrator complexes at distinct loci and clarify the interplay between IntS11 endonuclease activity and other Integrator subunits. For example, the non-catalytic subunits may be critical for the formation and targeting of the complex to specific loci and/or controlling RNAPII dynamics.

Finally, we note that the metazoan Integrator complex has parallels with the yeast Nrd1-Nab3-Sen1 (NNS) complex that (i) terminates transcription at both mRNA and snRNA loci and (ii) interacts with the RNA exosome (reviewed in Arndt and Reines 2015). Interestingly, the underlying molecular mechanisms of transcription termination carried out by these two complexes are quite distinct. NNS uses the Sen1 helicase to pull the nascent transcript out of the RNAPII active site, while Integrator likely promotes termination by taking advantage of its RNA endonuclease activity and providing an entry site for a 5’-3’ exonuclease (Connelly and Manley 1988; Eaton et al. 2018). There is currently conflicting data on whether the canonical “torpedo” exonuclease Rat1/Xrn2 is involved in termination at snRNA genes as only subtle termination defects have been observed at these loci when Rat1/Xrn2 is depleted from cells (O’Reilly et al. 2014; Fong et al. 2015; Eaton et al. 2018). Notably, Cpsf73 has been shown to behave as both an endonuclease and exonuclease (Yang et al. 2009), raising the possibility that IntS11 could support a ‘Rat1/Xrn2-like’ function and mediate termination. Future studies that compare and contrast the Integrator and NNS complexes, especially how their recruitment and termination activities are controlled, will shed light on this important facet of gene regulation. In summary, transcription attenuation through premature termination was first described decades ago in bacteria (reviewed in Naville and Gautheret 2010), and our work indicates that the metazoan Integrator complex can function analogously to limit expression from protein-coding genes.

## MATERIALS AND METHODS

### *Drosophila* cell culture and plasmid transfections

*Drosophila* DL1 cells were cultured at 25°C in Schneider’s *Drosophila* medium (Thermo Fisher Scientific 21720024), supplemented with 10% (v/v) fetal bovine serum (HyClone SH30910.03), 1% (v/v) penicillin-streptomycin (Thermo Fisher Scientific 15140122), and 1% (v/v) L-glutamine (Thermo Fisher Scientific 35050061). *Drosophila* S2 cells were cultured at 25°C in SFX-Insect (HyClone SH30278.01).

To generate eGFP and nLuc reporter plasmids, the indicated sequences were inserted into pMtnA eGFP SV40 (Kramer et al. 2015). To generate Integrator expression plasmids, each *Drosophila* cDNA was cloned into a previously described pUB-3xFLAG vector (Chen et al. 2012). Cloning details for all constructs and how they were transfected using Effectene (Qiagen 301427) or Fugene HD (Promega E2311) are provided in the **Supplemental Material**.

### Genome-scale RNAi screen

Double-stranded RNAs (250 ng/well) were pre-spotted in 384-well plates (Ambion AM8500). DL1 cells stably maintaining “pMtnA eGFP MALAT1” were grown for one passage in the absence of hygromycin B. 15,000 cells were then seeded in each well of the 384-well plates in 10 µL of serum-free Schneider’s *Drosophila* media. After 1 h, complete media (with serum) was added (20 µL/well). Cells were grown for 3 days and then treated with 10 µL of media containing CuSO_4_ (500 µM final concentration, Fisher BioReagents BP346-500) for 6 h. Cells were fixed (5% formaldehyde, Fisher BioReagents BP531-500) and stained with Hoechst 33342 (Sigma B2261) to visualize nuclei. Four images per well (eGFP and Hoechst 33342 staining) were captured at 20x magnification using an automated microscope (ImageXpress Micro, Molecular Devices) and analyzed using MetaXpress software. “Mean stain integrated intensity” of eGFP and “total cell number” were calculated for each image, and the median and interquartile ranges (IQR) were used to calculate a z-score for each plate: (Mean stain integrated intensity-median)/(IQR*74). Wells with robust Z-scores ≥1.3 or ≤−1.3 were considered hits and Gene Ontology (GO) analysis was performed using Database for Annotation, Visualization and Integrated Discovery (DAVID), v6.8 (https://david.ncifcrf.gov/home.jsp) with standard parameters.

### RNAi

Double-stranded RNAs (dsRNAs) from the DRSC (*Drosophila* RNAi Screening Center) were generated by *in vitro* transcription (MEGAscript kit, Thermo Fisher Scientific AMB13345) of PCR templates containing the T7 promoter sequence on both ends. Primer sequences are provided in **Supplemental Table S4**. Complete details of how cells were seeded, transfected (if applicable), and processed are provided in the **Supplemental Material**.

### Analysis of RNA expression

Total RNA was isolated using Trizol (Thermo Fisher Scientific 15596018) as per the manufacturer’s instructions. To detect full-length mature mRNAs, Northern blots using NorthernMax reagents (Thermo Fisher Scientific) and oligonucleotide probes were performed as previously described (Tatomer et al. 2017). Small RNAs were separated by 8% denaturing polyacrylamide gel electrophoresis (National Diagnostics EC-833) and electroblotted/UV crosslinked to Hybond N+ membrane (GE Healthcare RPN303B). ULTRAhyb-oligo hybridization Buffer (Thermo Fisher Scientific AM8663) was used as per the manufacturer’s instructions. All oligonucleotide probe sequences are provided in **Supplemental Table S4**. Blots were viewed and quantified with the Typhoon 9500 scanner (GE Healthcare) and quantified using ImageQuant (GE Healthcare).

For analysis of transcript levels by RT-qPCR, cDNA was reversed transcribed from total RNA using TaqMan Reverse Transcription Reagents (Thermo Fisher Scientific N8080234) according to the manufacturer’s instructions. Random hexamers were used, except for **Supplemental Fig. S5A** where oligo(dT) primer was used. RT-qPCR was then carried out in triplicate using Power SYBR Green Master Mix (Thermo Fisher Scientific 4367659). All RT-qPCR primers are provided in **Supplemental Table S4.**

The 5’ cap status of the MtnA small RNAs was characterized using Cap-Clip Acid Pyrophosphatase (CellScript C-CC15011H) and Terminator 5’-phosphate-dependent exonuclease (Lucigen TER51020). Ligation-mediated RACE was used to determine the 3’ ends of the MtnA small RNAs and full details are provided in the **Supplemental Material**.

### Analysis of protein expression

Protein levels were measured using standard Western blotting or immunofluorescence protocols and the details are provided in the **Supplemental Material**. Antibodies used and their conditions are provided in **Supplemental Table S5**.

### Chromatin Immunoprecipitation

ChIP-qPCR was performed as described in the **Supplemental Material**. Antibodies used and their conditions are provided in **Supplemental Table S5**. Primers used are provided in **Supplemental Table S4**.

### IP/Mass Spectrometry Analysis

Nuclear extracts were prepared from S2 cell lines stably expressing FLAG-IntS1, FLAG-IntS5, and naïve S2 cells as a negative control. Purifications were conducted using ∼10 mg of nuclear extract (2 mL). Details for FLAG affinity purification, sample digestion, nanoLC MS/MS analysis and database searching are provide in the **Supplemental Material**.

### RNA-seq

DL1 cells were treated for 3 days with a control (βgal) dsRNA or a dsRNA to deplete IntS9. CuSO_4_ was added for the last 14 h. Total RNA was isolated from three biological replicates and 1 µg of each was treated with DNase I (Qiagen 79254) and purified (RNeasy MinElute Cleanup Kit, Qiagen 74204). RNA quality was confirmed with a TapeStation (Agilent). RNA-seq libraries were generated using the TruSeq Stranded Total RNA preparation kit (Illumina RS-122-2302) and sequenced using a single-end 75 bp cycle run on an Illumina NextSeq 500. Details for how reads were mapped and analyzed are provided in the **Supplemental Material.** All RNA-seq datasets generated in this study are available for download from GEO (GSE136150). Data can also be viewed on the UCSC browser using the following link: https://genome.ucsc.edu/s/meishengxiao/dm6_DL1_Bgal_IntS9_WiluszLab_Upenn.

### Quantification and Statistical Analysis

For Northern blots, RT-qPCRs, and automated microscopy experiments, statistical significance for comparisons of means was assessed by Student’s t test. Unless otherwise indicated, the comparison was to the control (βgal dsRNA) treated samples. Statistical details and error bars are defined in each figure legend.

## ACKNOWLEDGMENTS

We thank Todd Albrecht for his help in generating *Drosophila* Integrator antibodies, the Andrulis lab for the Rrp40 antibody, William K. Russel and the UTMB Proteomics Core, the University of Pennsylvania High-Throughput Screening Core, Dan Beiting and Ana Misic for help with RNA-seq library preparation, as well as Karen Adelman and other members of the Cherry, Wagner, and Wilusz labs for helpful discussions. Supported by National Institutes of Health grants R35-GM119735 (to J.E.W.), K99-GM131028 (to D.C.T.), R01-AI122749 (to S.C.), R01-AI074951 (to S.C.), R01-GM134539 (to E.J.W.), and Welch Foundation grant H-1889 (to E.J.W.). S.C. is a recipient of the Burroughs Wellcome Investigators in the Pathogenesis of Infectious Disease Award. J.E.W. is a Rita Allen Foundation Scholar.

## AUTHOR CONTRIBUTIONS

D.C.T., S.C., and J.E.W. conceived the project and designed/analyzed the RNAi screen. D.C.T., N.D.E., D.L., J.Z.J., M.J., and K.-L.H. performed experiments and analyzed data. M.-S.X. analyzed the RNA-seq results. D.C.T., E.J.W., S.C., and J.E.W. wrote the manuscript with input from the other authors.

