## Supplemental Text & Figures for "The Integrator complex cleaves nascent mRNAs to attenuate transcription"

**Table of Contents**

**SUPPLEMENTAL METHODS**

Expression plasmid construction

Generation of stable cell lines

RNAi

Sequential RNAi / plasmid transfection experiments

Characterization of 5' end of MtnA small RNAs

Characterization of 3' ends of MtnA small RNAs using 3' ligation-mediated RACE

Analysis of protein expression by Western blotting and immunofluorescence

IntS12 antibody production

Chromatin immunoprecipitation (ChIP)-qPCR

IP/Mass spectrometry analysis

RNA-seq analysis

**SUPPLEMENTAL FIGURES**

**SUPPLEMENTAL FIGURE LEGENDS**

**SUPPLEMENTAL TABLE LEGENDS**

**SUPPLEMENTAL REFERENCES**

#### SUPPLEMENTAL METHODS

##### Expression plasmid construction

The intronless eGFP reporter driven by the MtnA promoter and terminating in the SV40 polyadenylation signal (**pMtnA eGFP SV40**) was previously described (Kramer et al. 2015) (https://www.addgene.org/69911/). The non-polyadenylated eGFP reporter (**pMtnA eGFP MALAT1**) was generated by replacing the SV40 polyadenylation signal with the mouse MALAT1 triple helix (Wilusz et al. 2012) followed by a hammerhead ribozyme (Dower et al. 2004). Specifically, the following sequence (triple helix in lowercase, hammerhead ribozyme in uppercase) was inserted between the NotI and SpeI sites:

```
gattcgtcagtagggttgtaaaggtttttcttttcctgagaaaacaaccttttgttttctcaggttttgctttttggcctttccct
agctttaaaaaaaaaaaagcaaaaCCTGTCACCGGATGTGTTTTCCGGTCTGATGAGTCCGTGAGGACGAAACAGG
```

To generate **pMtnA nLuc SV40**, pMtnA eGFP SV40 was cut with EcoRV and NotI and the eGFP ORF was replaced with the nano-Luciferase (nLuc) ORF:

```
ATGGTCTTCACACTCGAAGATTTCGTTGGGGACTGGCGACAGACAGCCGGCTACAACCTGGACCAAGTCCTTGAACAGGGAGGTGT
GTCCAGTTTGTTTCAGAATCTCGGGGTGTCCGTAACTCCGATCCAAAGGATTGTCCTGAGCGGTGAAAATGGGCTGAAGATCGACA
TCCATGTCATCATCCCGTATGAAGGTCTGAGCGGCGACCAAATGGGCCAGATCGAAAAAATTTTTAAGGTGGTGTACCCTGTGGAT
GATCATCACTTTAAGGTGATCCTGCACTATGGCACACTGGTAATCGACGGGGTTACGCCGAACATGATCGACTATTTCGGACGGCC
GTATGAAGGCATCGCCGTGTTCGACGGCAAAAAGATCACTGTAACAGGGACCCTGTGGAACGGCAACAAAATTATCGACGAGCGCC
TGATCAACCCCGACGGCTCCCTGCTGTTCCGAGTAACCATCAACGGAGTGACCGGCTGGCGGCTGTGCGAACGCATTCTGGCGGAC
TACAAAGACCATGACGGTGATTATAAAGATCATGACATCGATTACAAGGATGACGATGACAAGTAA
```

To generate eGFP reporter plasmids driven by the Integrator regulated promoters, a PacI site was first introduced by site-directed mutagenesis upstream of the MtnA promoter of pMtnA eGFP SV40. **pPepck1 eGFP SV40** was then generated from pMtnA eGFP SV40 by cutting with PacI and XhoI and replacing the MtnA promoter with the following sequence from the Pepck1 locus:

```
aaaaacagagccatatttacgtgcaactttcccgactgcttctgttgaactttttcttcgagtacccagaggattctgggtcttct
ggtaaactggattcggggtacaaagtcacaccaccggcagcaccttcatttctccttccccaaagacaaaaacaagagtggccaca
tgacaaccatccgggccatgctattcttggtgcgcccccagagtcacatgaatgtgaatactggcctccaaggcggccatggccaa
ttggggaggcccccggctcatctggatgccgcgagggggcggattagcatacaaattttccgtcaccgattctacaaataagcga
```

```
aagtttttttcgatttggcaaatgtccaaaaacgagaggccaacacatgtggtcaaacaggcgtaagtgggcgggcgcgtgtgatg
tggcccaccattcgatgttgcaccaatgtagctatatttgtggccaatgcagctatagacacatccagaatttacccagctgcagg
ggacaacaactgctgttgcctcagctgtccgtgaactggtttctatattcgcagcggccacatttgtatcgcataaaatcataaat
agaataattaaattaacagcgcgaaagtggcaaatatcagggtgttcaaaataacaacaaacgcttatcaatgcaaacgcactaga
aaacgctctcagcgccacatgaaaacaacactctgctgttgccatgtggggacataccccatatacgtatctgtatgcccatgccc
cccattccgtagacccccccccccacccacaaccaactcatcgtagctgcgccgcctgtgaaaaaccaaaacaatcgccgagcct
ccgccggcagccaatcagctcgtccggagtttgaccctctcactcggcgaggcgatcgtaaaacccgatttgcgtggcgatcggcc
gacttgttctagtcgcctcgatggccaggcgtcgccggctgggtataaaaggccccggatcggagcagttgGCATCAGTTACTCGT
GGCCAGAGTAA
```

**pHml eGFP SV40** was generated from pMtnA eGFP SV40 by cutting with PacI and XhoI and replacing the MtnA promoter with the following sequence from the Hml locus:

```
atttggtacccaaaagttatttctgtaggcaggttcaatacactttactaattttgacaccaatgcgccctctagcgagctacgag
acaagctgctgctcgaacgcatgaaccgtaaactgcttgcattcagcatttgcactgcttgaaccgataacttttaacggtcacaa
tgtagtacgttgaatatttgaaattagttcacccttggccaacattaaagtgattatattccaccacccaagttcaatgccattgg
agaagtttctgcgtaggacgctttacatccagcagctcttcgttgaccatataacttttgaatggcgacatatcagcagagggggc
taccccaagtcagaaaattcttacacccatcgtggactctagtaaaattatgacataggtcccggaaatgccgctatcaggaaggc
attagcatcagcttaaacttggcacagatttgttacattcgctttgctaatttaagaacggcgatgccgataagctgatgatggtg
ctttttttgcggatcgtggataaatagtgggcttgagcctaacaattccgagtgttctgataaaccagctacggaatcggcctat
aaaaggggctccccattcatggggaacattagtcgcgatcgcgcatcgaaatcgtcaagttcagtggatataattacagtgaaaaa
gtgctaagtgtaaaagtttttaaaagtgagttctaagctgga
```

**pCG8620 eGFP SV40** was generated from pMtnA eGFP SV40 by cutting with PacI and XhoI and replacing the MtnA promoter with the following sequence from the CG8620 locus:

```
ataatgtgctaaacctgatttatgcaattattaagatcccaaaacattttcaaaaaattgtgtaaatttcttaatactttatattc
ttaaaattttcgatgatttaaaacgataaaataattgttttaaaattacacacttggacgaaattcaaaattaaaattatatcaaa
ataatgtcatatttcacaaaatcgtaaaaaataataaaaggtatgcttataaaaatacgcagaatatttataatgcatataatatg
cagaaatacaaaagtaagaagaatttcgcaaaaatatgaaatatgtttgaggttacgttcggcaatgaaattatgtaacattaaaa
acacgtgatcaatggaagtaatttcccaaaaacccatgaatgcgatttggttaaaaatagctcagatgacgtcacatttatcgtgt
ggtttgtatgagtatgtgactgataaagaaaggcaaatgtttacagtccgaaagtcgcgattagcttggcgccaaagttgcctata
aaagctcatcgaccgaatgccgcagaagcCAGTCAAATTCAAGTGTTCAAGTGAAGCAGTCGAAACATAAGCAACAGATTCCGCAG
CAGTTATATCCAGTTCAGAGAAACAGTAGTTAATCAGTAATTAAAGTGTTGAGTGAATTTCTTATCGGGTCAGTTGTAAACTAGTG
CAAAA
```

**pSirup eGFP SV40** was generated from pMtnA eGFP SV40 by cutting with PacI and XhoI and replacing the MtnA promoter with the following sequence from the Sirup locus:

```
aactccagagtaatttcagtagccctttacgatttcggcttcttaacttatttggttgataagccaagataatgtccgggacccaa
ttcctaatcgcctgctctcaattaccgatgggactttgcgataagaacagggcgactgggcgaccttatagccgtgcaactaacct
ttgatataaaatgaaaagggtgagctcgctggatttcaacgatcgatggctctaatcagctttacatgtctttaatatgacaacat
aatgaaaagcatatagaacattacaaactccagataggcgcgattatacgtcggtttcttatacaaaaaaaaaaaaaaaaactgat
aaaaaatgccagttgataatatattaagaaatggtttgaacattttttgaatttaattattatgcagaagtgagcaaacaaatttgc
aatagattttaagaagtgtgatagtactataccattaaggcgtaataaaaaataaatgcaactaactgttaaatgtcttgagtaaa
tggttgaatgcacttaaaataaataaagttgcaatatttttcgcagtttctttctttcaataacggatttaaatttaaaagccagc
gttgccagatagcatgtagctagagttacagctcaatgcctagcacagtcatgcaccctggtgtgcgtctccgctgtgcgtattgt
atggctatatttagtacattttcagtgccACCACAACATATGGCACAATAAATAAAGCAACTAAACAAGCAACGCGGGCTATAAAA
GCGAAACGCAGCCTCGCAGATTGCAATTCAGACGCGAAGCAACAAGCAAGTGACATTGACTCCCAGCCGAAAGTGAGACTTTTGAG
TAAACACACAAAGTGTAACTGCGACTAAGAAGCGAACAAAAACGAAGATTCGCCGGCATCATCTTGCATAACCGATAAAGGAAATC
CAAACAAAAAACAAA
```

**ppst eGFP SV40** was generated from pMtnA eGFP SV40 by cutting with PacI and XhoI and replacing the MtnA promoter with the following sequence from the pst locus:

```
CTCCGACAGCGACTATATGTATTTTGTATTTTGTCTGGCACTGCCTTTTGGCTCCGTTCCGTTCCGTTCCTTTCTCTGCCGCCGCT
TTCCCCTGCACTTTTCCTCCGGCAGATCCAGCTGATGTTGCTGTCGCGTTGGAGTTATCGCTACAATCGCTGCGCCCCGTGGATTC
TCGATCGACACGCGGCGATCTGCGAGATTCGAATGGGGCAGGAGCTGGTGTTGTTGCTGCTGCTGCTGCAAGATATCGCGAGATTG
GCACGCCGCGAACTTAAAAAACTCTGTATTTACACCGACGCATGAGACTGACACGTAGGGTTGTTTGGCTCGTGCTGGAACCGATA
GGCgtagctggagcggagccggttcgattgcacggctatcgataggcgggaggcagcattcatagtttagggttgccaaagctggc
agtactaagtttagaatctgatttttaggtcaccaagtactaaaaatgtttgcaagcaaaacttttgtcagcaagttttcaatgt
aaatgtatatttttaaatggatttgcctatatatatatatatatatatttttttttttttataatataatatgtgctcgctagaaat
acaaagcaaatatctgattagcatgagctcatcgtctcgttaCATTTTGGGTTTAGCACGCTAGCAGCTTTTTAAATTTTGCTCAC
CAAAGAAGGTGACGGTGGAATACGGGGTATTCATATAAAGGTATGATCCCTAGAGAGCTTTTCTGCCGTCTTGCCTTTTCGGTATT
TCGAGTACGATCGTTCAATTAGTGTGCGCCTTAACTTAAAAGCTGGTGATTCCGTGGCACAACTCCATTAGCAGCACAACTGTGGC
AACGGAAAGGTGACAACTTAGGG
```

**pCG6770 eGFP SV40** was generated from pMtnA eGFP SV40 by cutting with PacI and XhoI and replacing the MtnA promoter with the following sequence from the CG6770 locus:

```
GAATAAAACGGAAGGTTACCCCATTCATAACTTCTCAGAGAAAGAGATTTGTGAGGCAAAGGTCAATAGTAAATATTTAGAATAAC
TATGTTGAAAAGAGACTGTATCAGCAAGTTTAGAACATTTGCTATATTTACAAAATGAGACGTGGGATCTCTCGTTCTGAGTTCTA
GTACAACAAAATTGTATCTAGAATATTCACGGAAACCCAGTTTCTTTTTAAAGCCATTAAAAAAGAACCATCGAGAAATCTAAAGA
TTGATTTGCTACCAACGAATTGTAAATAAAGTGGTATATTGTGGCAGGTTCTTGAAACAACATTGTTGAGGCCGTTCTATCTTATC
TAAGATGGTGACTACGATCTATTGCGTCGTTTGAAGACGAACTTTCAGAACGAATACCTGCCAAATGACTAACTAGAGCGATAGTT
AGCACCTTATCAACGCCAGAAGGCTATGTATATATGTATGTACGTAACATTTGTTTATATAAAATCCATTAAACCAAAACAAACTG
AAAGCTTTTCGCTATTCAGAAATAAACAAATGCGTTGCCTGGTGGGAGGGAAGTGGGTCAACCGATCAGGTTCTAATGCACCAAAA
TAAATATCCATTATGAGTATATCTTTTGTAAATCAAAAATGTTATTCGAATTATAATTATTGCTAAATTGTTTCTTGTGCCCTaac
attcaaatattgttgagtgtaaaaaaatAAACAAATTTTGAGCAGTGTATCCAGAACATATGATGCATTTAATTCTATATTTATTG
GGTGGAGAAACAAATGCAGGGTAAATGCACTTTGATGGTCGTTTGCGGAGGGCCAAGGTGGGTGTGTGTTATGTAACCACTCAGGA
ACGTGTGTATGCTTAAAAGTTCTTAAATTATTAAAATAAATGAACAAGGCGAAATAATATGTTTGGCGATTTCAGTTTTTTTGTTT
ACCGCGCActttgtaaacagcaaaaaattaaattgcgcagcaaaaattgtcaaggaggcactctcttgagcgcaaaagggttgtca
ggcccaataactatgaattttaataactaaaaattcttttacataaaaaagagcaaatatattttgtaaaagttcagtagaaagtt
atagaaaaattaaatatttttagtaactacacgtaagcaacaattaaatgcccttctagtagtttctttagctaaatccggcaact
ctgacggagagagagctttggttacggaacagaaagcttgcgcgcctataaaagccgctgctcgaccgaaaaaATTCAATTCCGTT
CGCAGCTCAGCGAAAAGTGAACAAAAAAAAGCAGAAGAGAAATTTGGAAAGCAGCCTGCAAGTTTGCAGAGAAACAAAAAGCAGAC
ACCAACCAAGAACGCAGTTAGTTTTGCAACCTTAAATCGCCATT
```

**pUbi-p63E eGFP SV40** was generated from pMtnA eGFP SV40 by cutting with PacI and EcoRV and replacing the MtnA promoter with the following sequence from the Ubi-p63E locus:

```
taaGATCTTGTCGCCGGAACGCAGCGACAGGGATTCCAATGTGTCCGTATCCTTTAGGCTTTTGCCCTTCAGTTCCAGACGAAGCG
ACTGGCGATTCGCGTGTGGGGTCTGCTTCAGGGTCTTGTGAATTAGGGCGCGCAGATCGCCGATGGGCGTGGCGCCGGAGGGCACC
TTCACCTTGCCGTACGGCTTGCTGTTCTTCGCGTTCAAAATCTCCAGCTCCATTTTGCTTTCGGTGCGCTTGCAATCAGTACTGTC
CAAAATCGAAAATCGCcgaaccgtagtgtgaccgtgcggggctctgcgaaaataaactttttttaggtatatggccacacacgggga
aagcacagtggattatatgtattaatatattatccaggttttcattacttatccagatgtaagcccacttaaagcgatttaacaat
tatttgccgaaagagtataaacaaattttacttaaaaatggattaagaaaagcttgtgtaagattatgcgcagcgttgccagatag
ctccatttaaaacacttcaaaaacaataagttttgaaaatatatacataaatagcagtcgttgccGCAACGCTCAACACATCACAC
TTTTAAAACACCCTTTACCTACACAGAATTACTTTTTAAATTTCCAGTCAAGCTGCGAGTTTCAAAATTATAGCCGGTAGAGAAGA
CAGTGCTATTTCAAAAGCAAACTAACAAGGGTCTTAAATTCCAAAACACCAATCCTAACAAGCCTTGGACTTTTgtaagtttagat
caaaggtggcattgcattcaatgtcatggtaagaagtaggtcgtctaggtagaaatcctcattcagccggtCAAGTCAGTACGAGA
AAGGTCTCAATTTGAAATTGTCTTAAAAATATTTTATTGTTTTGTACTGTGgtgagtttaaacgaaaaacacaaaaaaaaaagtgat
acatagaaatcataaaaaattttaatacaaggtattcgtacgtatcaaaaacatttcggcacaatttttttttctctgtactaaagt
gttacgaacactacggtatttttttagTGATTTTCAACGGACACCGAAGGTATATAAACAGCGTTCGCGAACGGTCGCCTTCAAAAC
```

CAATTGACATTTGCAGCAGCAAGTACAAGCAGAAAGTAAAGCGCAATCAGCGAAAAATTTATACTTAATTGTTGGTGATTAAAGTA
CAATTAAAAGAACATTCTCGAAAGTCACAAGAAACgtaagtttttaactcgctgttaccaattagtaataagagcaacaagacgtt
gagtaatttcaagaaaaactgcatttcaaggtctttgttcggccatttttttttattcaacgctctacgtaattacaaaataagaa
attggcagccacgcatctcgttttcccaatgaattggcatcaaaacgcaaacaaatctataaataaaacttacgtgttgattttcg
ccaagatttattggcaaattgtgaaattcgcagtgacgtatttgaaaattcgagaaatcacatacgcactcgagcatttgtgtgca
tgttattagttagttgtttagttaattgaagtattttaccaacgaaatccacttatttttagctgaaatagagtaggttgcttaaa
caaagccacgtctgaaaatttcttattgcttgtagttgtgacgtcaccatatacacacaaaataatgtgtatgcatgcattTCAGC
TGTGTATACATACATGCACACACTCGCAACACGAAAACGATGACGAAGCAACGGAACAAAGGTTTCTCAACTACCCTTTGTTCCCT
GTTTCTTCGCTTTCCTTTGTTCCAATATTCGTAGAGGGTTAATAGGGGTTTCTCAACAAAGTTGGCGTCGATAAATAAGTTTCCCA
TTTTTATTCCCCAGCCAGAAAgttagtttcaatagttttgtaatttcaacgaaactcgtctgatttcgtactaattttccacatct
ctattttcttcccgcagaataatccatctagagat

To generate the **pMtnA UTR 1-55 eGFP SV40**, **pMtnA UTR Δ41-55 eGFP SV40**, and **pMtnA UTR Δ21-55 eGFP SV40** plasmids, the following sequences were inserted between the PacI and XhoI sites:

<u>pMtnA UTR 1-55</u>
ttgcaggacaggatgtggtgcccgatgtgactagctctttgctgcaggccgtcctatcctctggttccgataagagacccagaact
ccggcccccccaccgcccaccgccacccccatacatatgtggtacgcaagtaagagtgcctgcgcatgccccatgtgccccaccaag
agctttgcatcccatacaagtccccaaagtggagaaccgaaccaattcttcgcgggcagaacaaaagcttctgcacacgtctccac
tcgaatttggagccggccggcgtgtgcaaaagaggtgaatcgaacgaaagacccgtgtgtaaagccgcgtttccaaaatgtataaa
accgagagcatctggccaatgTGCATCAGTTGTGGTCAGCAGCAAAATCAAGTGAATCATCTCAGTGCAACTAAAG

<u>pMtnA UTR Δ41-55</u>
ttgcaggacaggatgtggtgcccgatgtgactagctctttgctgcaggccgtcctatcctctggttccgataagagacccagaact
ccggcccccccaccgcccaccgccacccccatacatatgtggtacgcaagtaagagtgcctgcgcatgccccatgtgccccaccaag
agctttgcatcccatacaagtccccaaagtggagaaccgaaccaattcttcgcgggcagaacaaaagcttctgcacacgtctccac
tcgaatttggagccggccggcgtgtgcaaaagaggtgaatcgaacgaaagacccgtgtgtaaagccgcgtttccaaaatgtataaa
accgagagcatctggccaatgTGCATCAGTTGTGGTCAGCAGCAAAATCAAGTGAATCATC

<u>pMtnA UTR Δ21-55</u>
ttgcaggacaggatgtggtgcccgatgtgactagctctttgctgcaggccgtcctatcctctggttccgataagagacccagaact
ccggcccccccaccgcccaccgccacccccatacatatgtggtacgcaagtaagagtgcctgcgcatgccccatgtgccccaccaag
agctttgcatcccatacaagtccccaaagtggagaaccgaaccaattcttcgcgggcagaacaaaagcttctgcacacgtctccac
tcgaatttggagccggccggcgtgtgcaaaagaggtgaatcgaacgaaagacccgtgtgtaaagccgcgtttccaaaatgtataaa
accgagagcatctggccaatgTGCATCAGTTGTGGTCAGCA

To generate the IntS1, IntS5, and IntS11 expression plasmids, each respective *Drosophila* cDNA was cloned into a previously described pUB-3xFLAG vector (Chen et al. 2012). PCR primers are provided in **Supplemental Table S4**. The IntS11 E203Q mutation (GAG to CAG) was subsequently introduced using site-directed mutagenesis. All plasmids were sequenced to confirm identity.

**Generation of stable cell lines**

To generate DL1 cells stably maintaining the eGFP reporters, 2 x $10^6$ cells were first plated in complete media in 6-well dishes. After 24 h, 2 µg of pMtnA eGFP SV40, pMtnA eGFP MALAT1, pMtnA UTR 1-55, pMtnA UTR Δ41-55, or pMtnA UTR Δ21-55 plasmid was transfected using Effectene (Qiagen 301427; 16 µL Enhancer and 30 µL Effectene Reagent). On the following day, 150 µg/mL hygromycin B was added to the media to select and maintain the cell population.

To generate DL1 cells stably maintaining the Flag-tagged IntS11 transgenes, 2 x $10^6$ cells were first plated in complete media in 6-well dishes. After 24 h, 1.8 µg of pUB IntS11 WT or pUB IntS11 E203Q along with 180 ng of pAct Puro [puromycin resistance gene driven by the actin promoter; obtained from Sara Cherry] were transfected using Effectene (Qiagen 301427; 16 µL Enhancer and 30 µL Effectene Reagent). On the following day, 0.5 µg/mL puromycin was added to the media to select and maintain the cell population.

To generate S2 cells stably maintaining the Flag-tagged IntS1 or IntS5 transgenes, 2 x $10^6$ cells were first plated in complete media in 6-well dishes. After 24 h, 2 µg of pUB Flag-IntS1 or Flag-IntS5 along with 50 ng of pIZ CoBlast [balsticidin resistance gene driven by the copia promoter] were transfected using Fugene HD (Promega E2311). On the following day, 10 µg/mL blasticidin was added to the media to select and maintain the cell population.

**RNAi**

Double-stranded RNAs from the DRSC (*Drosophila* RNAi Screening Center) were generated by *in vitro* transcription (MEGAscript kit, Thermo Fisher Scientific AMB13345) of PCR templates containing the T7 promoter sequence on both ends. Primer sequences are provided in **Supplemental Table S4**. Knockdown experiments in 12-well dishes were then performed by bathing 500,000 cells with 2 µg of dsRNA. Cells were incubated for 3 days and, if applicable, a final concentration of 500 μM $CuSO_4$ (Fisher BioReagents BP346-500) or 50 μM $CdCl_2$ (Sigma C3141) was added for the final indicated times.

For the validation experiments shown in **Supplemental Fig. S2A-C**, cells were resuspended after 4 h of $CuSO_4$ treatment and 10% of the sample was plated (50 μl/well, 3 wells per dsRNA) in 96-well plates (Corning 3904). After an additional 2 h (resulting in a total of 6 h $CuSO_4$ treatment), cells were fixed in 5% formaldehyde (Fisher BioReagents BP531-500) and counterstained with Hoechst 33342 (Sigma B2261). Four images per well (eGFP and Hoechst 333342) were captured at 20x magnification using an automated microscope (ImageXpress Micro, Molecular Devices) and analyzed with MetaXpress software to quantify the integrated eGFP intensity and total cell counts. Averages for each well were calculated and normalized to control (treatment with βgal dsRNA) cells.

**Sequential RNAi / plasmid transfection experiments**

To confirm that Integrator acts on the MtnA promoter regardless of the downstream ORF **(Supplemental Fig. S2F)**, 1 x $10^6$ DL1 cells were first bathed in 2 μg of dsRNA in 12-well dishes. After 24 h, Effectene (Qiagen 301427; 4 μL Enhancer and 5 μL Effectene Reagent) was used to transfect 500 ng "pMtnA nLuc SV40" plasmid. Cells were treated with 500 μM $CuSO_4$ for the final 14 h of the 72 h experiment. 2 h before collection, cells were resuspended and 100 μL of cells/well were plated in a 96 well dish. Cells were lysed in 100 μL of Glo Lysis Buffer (Promega E2661) for 10 min at room temperature. Luciferase assays were then performed by mixing equal volumes of cell lysate and Nano-Glo (Promega N1120) in black 96-well plates for 5 min. Luminescence was measured on a GloMax Explorer Multimode Plate Reader (Promega).

To determine if the endonuclease activity of IntS11 is required for inhibition of the "pMtnA eGFP SV40" reporter **(Supplemental Fig. S4)**, 3 x $10^6$ DL1 cells were first bathed in 4 μg of dsRNA in 6-well dishes. After 24 h, Effectene (Qiagen 301427; 16 μL Enhancer and 30 μL Effectene Reagent) was used to transfect 1 μg "pMtnA eGFP SV40" plasmid and 100 ng of either "pUB IntS11 WT," "pUB IntS11 E203Q," or an empty pUB vector with no inserted ORF. Transfected cells were incubated for 42 h, then resuspended in media containing 500 μM $CuSO_4$ and transferred to new 6 well dishes. After 4 h of $CuSO_4$ treatment, cells were again resuspended and 10% of the sample was plated (50 μl/well, 3 wells per dsRNA) in 96-well plates (Corning 3904). After 2 h (resulting in a total of 6 h $CuSO_4$ treatment), cells were fixed in 5% formaldehyde (Fisher BioReagents BP531-500) and processed for FLAG immunofluorescence (see below) or quantification of eGFP levels. Four images per well (eGFP, TexasRed, and Hoechst 333342) were captured at 20x magnification using an automated microscope (ImageXpress Micro, Molecular Devices) and analyzed with MetaXpress software to measure integrated intensity and total cell number. Averages for each set of images were calculated and normalized to control (βgal) dsRNA treated cells.

To determine whether eGFP reporters driven by the indicated promoters are regulated by Integrator **(Fig. 6)**, $1 \times 10^6$ DL1 cells were first bathed in 2 μg of dsRNA in 12-well dishes. After 24 h, Effectene (Qiagen 301427; 4 μL Enhancer and 5 μL Effectene Reagent) was used to transfect 500 ng of each reporter plasmid. Total RNA was extracted after an additional 48 h incubation.

**Characterization of 5' end of MtnA small RNAs**

The 5' cap status of the MtnA small RNAs was determined by incubating 20 μg of total RNA with Cap-Clip Acid Pyrophosphatase (CellScript C-CC15011H) or buffer alone for 90 min at 37°C. After extracting with acid-phenol:chloroform (Thermo Fisher Scientific AM9720) and ethanol precipitation, samples were split in half and incubated with Terminator 5'-phosphatedependent exonuclease (Lucigen TER51020) or buffer alone for 90 min at 30°C. Reactions were then extracted with acid-phenol:chloroform, ethanol precipitated, and analyzed using 8% denaturing polyacrylamide Northern gels.

**Characterization of 3' ends of MtnA small RNAs using 3' ligation-mediated RACE**

DL1 cells were treated with Mtr4 dsRNAs for 3 days and $CuSO_4$ was added for the last 14 h. Total RNA was isolated using Trizol and 2 μg was ligated to 10 pmol of the 3' RNA adapter oligo (Sigma) **(Supplemental Table S4)** using T4 RNA Ligase I (NEB M0204S) following the manufacturer's protocol at 20°C for 6 h. The reaction was acid-phenol:chloroform extracted (Thermo Fisher Scientific AM9720) and ethanol precipitated. Reverse transcription (RT) was performed using M-MLV (Thermo Fisher Scientific 28025013) using the 3' RACE RT Primer. 2 μL of cDNA was used as a template for 20 PCR cycles using PFU and a gene-specific MtnA 5' forward primer and the RT Primer (95°C melting 15 s, 60°C annealing 15 s, 72°C extension 30 s). 2 μL of this PCR was then added to a new reaction for an additional 20 PCR cycles using the nested forward and RT primers **(Supplemental Table S4)**. The resultant PCR products were cloned using the Zero Blunt TOPO PCR Cloning Kit (Thermo Fisher Scientific) and sequenced.

**Analysis of protein expression by Western blotting and immunofluorescence**

For Western blotting, cells were first resuspended in PBS, pelleted at 2,100 rpm for 5 min, and then resuspended in RIPA buffer (150 mM NaCl, 1% Triton X-100, 50 mM Tris pH 7.5, 0.1% SDS, 0.5% sodium-deoxycholate, and protease inhibitors [Roche 11836170001]). Lysates were passed 10 times through a 28.5 gauge needle and cleared by centrifugation at 12,700 rpm for 20 min at 4°C. Lysates were then resolved on a NuPAGE 4-12% Bis-Tris gel

(Thermo Fisher Scientific NP0323) and transferred to a PVDF membrane (Bio-Rad 1620177). Antibody incubation conditions are summarized in **Supplemental Table S5**. Membranes were processed using SuperSignal West Pico Chemiluminescent Substrate (Thermo Fisher Scientific PI34080).

For immunofluorescence, DL1 cells were fixed in 5% formaldehyde (Fisher BioReagents BP531-500), washed 3 times in PBS for 10 min, incubated at room temperature for 90 min in blocking buffer (2% BSA in PBST [PBS + 0.1% Triton-X100]), and then incubated with anti-FLAG antibody **(Supplemental Table S5)** overnight at 4°C. After 3 washes (each 10 min) in PBST, cells were incubated simultaneously in Hoechst 33342 (Sigma B2261) and AlexaFluor594 goat anti-mouse IgG secondary antibody **(Supplemental Table S5)** for 60 min at room temperature. After 3 washes (each 10 min) in PBST, signal was captured by automated microscopy, as described above.

**IntS12 antibody production**

Full-length *Drosophila* Ints12 was cloned into pET-49b (Novagen) and transformed into BL21 competent cells. 100 mL of an overnight culture was expanded into 2 L of LB broth and incubated at 30°C with shaking until $OD_{600}$ reached 0.6-0.7. IPTG (Sigma) was then added to a final concentration of 0.4 mM for 4 h to induce protein expression. Cells were pelleted from 1 L of LB and resuspended in 50 mL cold lysis buffer (1X PBS, 2 μg/mL Aprotinin, 2 μg/mL Leupeptin, 2 μg/mL Pepstatin, 0.5 mM DTT, 1 mM PMSF, 1% Triton X-100, 1 mg/mL lysozyme, 5 μg/mL DNase I, and 10 mM Imidazole [pH 8]) and rotated for 1 h at 4°C. Sonication (10 cycles of 10 sec pulsing/30 sec resting on ice) was carried out to increase yield before spinning at 20,000 *g* at 4°C for 30 min to remove debris. Cleared lysate was mixed with 1 mL of pre-washed (3 cycles of cold PBS) Ni-NTA agarose slurry (Qiagen) and rotated overnight at 4°C. Agarose was washed (10 min rotation at 4°C each time) 3 times in 25 mL cold buffer I (200 mM NaCl, 20 mM Imidazole, 1% Triton X-100, 0.5 mM DTT, 1 mM PMSF, and 10% glycerol in PBS) and once in 25 mL cold buffer II (buffer I without Triton X-100). One additional wash with 1 mL cold buffer II was performed to transfer agarose to an eppendorf tube. IntS12 protein was eluted from agarose with 1 mL of elution buffer (250 mM Imidazole in PBS) by rotating for 10 min at 4°C (repeated 5 times). Eluted protein was then dialyzed overnight against 1 L of PBS in 3.5 KD MWCO membrane tubing (Spectrum Labs) and concentrated afterward in 10 KD cut-off Amicon Ultra-4 spin tube (Millipore) to 1/10$^{th}$ of original volume. 175 μg (1μg/μL) of IntS12 protein was used to raise an antibody in a guinea pig (Cocalico Biologicals, USA). To purify the antibody from crude serum, a SulfoLink Coupling Resin (Thermo) column was built as per the manufacturer's instructions. Briefly, 4 mL of resin slurry was packed into a 10 mL capacity chromatography column (Bio-Rad). Two full volume washes (Coupling buffer: 50 mM Tris [pH 8] and 5 mM EDTA [pH 8]) were performed to equilibrate the resin before adding 10 mL of diluted antigen (4 mg antigen in 50 mM Tris [pH 8], 5 mM EDTA [pH 8], and 25 mM TCEP [pH 8]). The mixture was incubated overnight at 4°C before washing the resin with coupling buffer, blocking nonspecific binding sites on the resin with coupling buffer containing 50 mM L-Cysteine-HCl for 30 min rotation at RT, washing the resin with 1 M NaCl, and PBS twice sequentially. Serum from the same guinea pig was pooled and loaded onto the column. The mixture was incubated overnight at 4°C before washing the resin with PBS 3 times and then eluting the antibody with 15 mL of 0.1 M glycine [pH 2.7]. Each 1 mL of eluted antibody solution was immediately neutralized by collecting it into a tube containing 60 μL of 1 M Tris [pH 8.8]. Sodium chloride (150 mM final) was added into the pooled solution before concentrating through a 10 KD cut-off Amicon Ultra-15 spin tube

(Millipore) to 1/5$^{th}$ of original volume. The concentration was adjusted to 2 μg/μL with 50 mM Tris [pH 7.5] before adding an equal amount of 100% glycerol and BSA to a final concentration of 100 μg/mL. Antibody specificity was then confirmed using immunoblotting.

**Chromatin immunoprecipitation (ChIP)-qPCR**

A 10-cm dish of 5 x $10^7$ DL1 cells was harvested into a 15 mL tube and centrifuged at 1,500 *g* for 2 min. Cells were then washed with 10 mL PBS and centrifuged at 1,500 *g* for 2 min. The cell pellet was resuspended in 10 mL of Fixing Buffer (50 mM Hepes pH 7.5, 100 mM NaCl, 1 mM EDTA pH 8.0, 0.5 mM EGTA pH 8.0 with 1% formaldehyde) and incubated at room temperature for 30 min. 0.5 mL of 2.5 M glycine was then added (final concentration of 0.125 M) and incubated at room temperature with rotation for 5 min, centrifuged at 1,500 *g* for 2 min, and washed two times with 10 mL PBS. Cells were lysed using lysis buffer (50 mM HEPES pH 7.9, 140 mM NaCl, 1 mM EDTA, 10% glycerol, 0.5% NP-40, 0.25% Triton X-100) for 10 min on ice and centrifuged at 1,500 *g* for 2 min. The pellet was then washed 2x in Wash Buffer (10 mM Tris-HCl pH 8.1, 200 mM NaCl, 1 mM EDTA pH 8.0, 0.5 mM EGTA pH 8.0) and resuspended in 1 mL Shearing Buffer (0.1% SDS, 1 mM EDTA, 10 mM Tris-HCl pH 8.1). The suspension was sonicated at 4°C using a Covaris S220 machine to obtain 500 bp DNA fragments in TC12x12 tubes with AFA fiber (Settings: Time- 15 min, Duty Cycle- 5%, Intensity- 4, Cycles per Burst- 200, Power mode Frequency- Sweeping, Degassing mode- Continuous, AFA Intensifier- none, Water level- 8). To the 1 mL of sheared chromatin, 115 μL of 10% Triton X-100 and 34 μL 5 M NaCl was added per mL of sheared chromatin, so that the final concentration of the sample is 1% Triton X-100 and 150 mM NaCl. Sheared chromatin was pre-cleared with protein A/G beads and 10 μL was reserved as input control. For each IP sample, 100 μL of sheared chromatin was diluted to 1 mL using IP Buffer (0.1% SDS, 1 mM EDTA, 10 mM Tris-

HCl pH 8.1, 1% Triton X-100, 150 mM NaCl) and incubated overnight at 4°C with 10 μL of serum. The next day, lysates were immunoprecipitated with protein A/G beads for 2 h at 4°C and washed once with low salt buffer (0.1% SDS, 1% Triton X-100, 2 mM EDTA, 20 mM Hepes pH 7.9, 150 mM NaCl), twice with high salt buffer (0.1% SDS, 1% Triton X-100, 2 mM EDTA, 20 mM Hepes pH 7.9, 500 mM NaCl), once with LiCl buffer (100 mM Tris-HCl pH 7.5. 0.5 M LiCl, 1% NP-40, 1% Sodium Deoxycholate), and once with TE. Immunocomplexes were eluted and de-crosslinked at 65°C overnight with Proteinase K and RNase A. DNA was extracted by phenol-chloroform and ethanol precipitated. DNA was resuspended in 100 uL, and 2 uL was used for each qPCR reaction.

**IP/Mass spectrometry analysis**

***FLAG affinity purification.*** Nuclear extracts were prepared from S2 cell lines stably expressing FLAG-IntS1, FLAG-IntS5, and naïve S2 cells as a negative control. Purifications were conducted using ~10 mg of nuclear extract (2 mL). To 2 mL of nuclear extract, 200 μL of a 50% slurry of anti-FLAG affinity resin (Sigma A2220) previously equilibrated in Buffer D [20 mM HEPES/KOH pH 7.9, 20% glycerol, 0.1 M KCl, 0.2 mM EDTA, 0.5 mM DTT] was added and rotated at 4°C for 3 h. Following the incubation, the beads were spun down at 3K at 4°C for 5 min. Beads were washed in 1 mL of Buffer D containing 150 mM NaCl with rotation for 5 minutes and then beads were spun down again. In total, the beads were washed four times. The final wash contained 50 mM $(NH_4)HCO_3$ to facilitate downstream analysis by mass spectrometry. In the case of Western blotting, the washed beads were incubated with 2x Laemmli Sample Buffer (Bio-Rad 161-0737) and resolved on 10% SDS-PAGE.

***Sample digestion.*** The proteins on beads were solubilized with 5% SDS, 50 mM TEAB pH 7.55 in a final volume of 25 μL. The sample was then centrifuged at 17,000 *g* for 10 min to remove any debris. Proteins were reduced by making the solution 20 mM TCEP (Thermo 77720) and incubated at 65°C for 30 min. The sample was then cooled to room temperature and 1 µL of 0.5 M iodoacetamide acid was added and allowed to react for 20 min in the dark. 2.75 µL of 12% phosphoric acid was added to the protein solution. 165 µL of binding buffer (90% Methanol, 100 mM TEAB final; pH 7.1) was then added to the solution. The resulting solution was added to an S-Trap spin column (protifi.com) and passed through the column using a bench top centrifuge (30 seconds at 4,000 *g*). The spin column was washed with 400 µL of binding buffer and centrifuged. This was repeated two more times. Trypsin was added to the protein mixture in a ratio of 1:25 in 50 mM TEAB, pH 8 and incubated at 37°C for 4 h. Peptides were eluted with 80 µL of 50 mM TEAB, followed by 80 µL of 0.2% formic acid, and finally 80 µL of 50% acetonitrile, 0.2% formic acid. The combined peptide solution was then dried in a SpeedVac concentrator and resuspended in 2% acetonitrile, 0.1% formic acid, 97.9% water and placed in an autosampler vial.

***NanoLC MS/MS analysis.*** Peptide mixtures were analyzed by nanoflow liquid chromatography-tandem mass spectrometry (nanoLC-MS/MS) using a nano-LC chromatography system (UltiMate 3000 RSLCnano, Dionex), coupled on-line to a Thermo Orbitrap Fusion mass spectrometer (Thermo Fisher Scientific, San Jose, CA) through a nanospray ion source (Thermo Scientific). A trap and elute method was used. The trap column was a C18 PepMap100 (300 µm X 5 mm, 5 µm particle size) from Thermo Scientific. The analytical column was an Acclaim PepMap 100 (75 µm X 25 cm) from Thermo Scientific. After equilibrating the column in 98% solvent A (0.1% formic acid in water) and 2% solvent B (0.1% formic acid in acetonitrile (ACN)), the samples (1 µL in solvent A) were injected onto the trap column and subsequently eluted (400 nL/min) by gradient elution onto the C18 column as follows: isocratic at 2% B, 0-5 min; 2% to 32% B, 5-100 min; 32% to 50% B, 100-108 min; 50% to 90% B, 108-109 min; isocratic at 90% B, 109-114 min; 90% to 2%, 114-115 min; and isocratic at 2% B, till 130 min.

All LC-MS/MS data were acquired using XCalibur, version 2.1.0 (Thermo Fisher Scientific) in positive ion mode using a top speed data-dependent acquisition (DDA) method with a 3 sec cycle time. The survey scans (*m/z* 350-1500) were acquired in the Orbitrap at 120,000 resolution (at *m/z* = 400) in profile mode, with a maximum injection time of 50 msec and an AGC target of 400,000 ions. The S-lens RF level was set to 60. Isolation was performed in the quadrupole with a 1.6 Da isolation window, and CID MS/MS acquisition was performed in profile mode using rapid scan rate with detection in the orbitrap (res: 35,000), with the following settings: parent threshold = 5,000; collision energy = 35%; maximum injection time 100 msec; AGC target 500,000 ions. Monoisotopic precursor selection (MIPS) and charge state filtering were on, with charge states 2-6 included. Dynamic exclusion was used to remove selected precursor ions, with a +/- 10 ppm mass tolerance, for 60 sec after acquisition of one MS/MS spectrum.

***Database searching.*** Tandem mass spectra were extracted and charge state deconvoluted by Proteome Discoverer (Thermo Fisher, version 1.4.1.14). Deisotoping was not performed. All MS/MS spectra were searched against a Uniprot *Drosophila* database (version 06-27-2018) using Sequest. Searches were performed with a parent ion tolerance of 5 ppm and a fragment ion tolerance of 0.60 Da. Trypsin was specified as the enzyme, allowing for two missed cleavages. Fixed modification of carbamidomethyl (C) and variable modifications of oxidation (M) and deamidation of asparagine and glutamine, were specified in Sequest. Scaffold (version Scaffold_4.8.7, Proteome Software Inc., Portland, OR) was used to validate MS/MS based peptide and protein identifications. Peptide identifications were accepted if they could be established at greater than 95% probability. Peptide probabilities from X! Tandem and Sequest were assigned by the Scaffold Local FDR algorithm. Peptide Probabilities were assigned by the Peptide Prophet algorithm (Keller et al. 2002) with Scaffold delta-mass correction. Protein identifications were accepted if they could be established at greater than 95% probability and contained at least 2 identified peptides. Protein probabilities were assigned by the Protein Prophet algorithm (Nesvizhskii et al. 2003). Proteins that contained similar peptides and could not be differentiated based on MS/MS analysis alone were grouped to satisfy the principles of parsimony.

**RNA-seq analysis**

Sequencing reads were filtered to remove low quality bases and adaptor sequences using fastp with default parameters (Chen et al. 2018) and then mapped to the dm6 genome (BDGP6.22, downloaded from Ensembl) using TopHat2 (version 2.0.14) with default parameters (Kim et al. 2013).

| **Sample** | **Total reads** | **Mappable reads (% of total)** |
|---|---|---|
| β-gal dsRNA Rep1 | 39,746,517 | 37,038,547 (93.2%) |
| β-gal dsRNA Rep2 | 45,567,267 | 43,137,125 (94.7%) |
| β-gal dsRNA Rep3 | 45,494,677 | 41,958,788 (92.2%) |
| IntS9 dsRNA Rep1 | 35,171,350 | 33,324,630 (94.7%) |
| IntS9 dsRNA Rep2 | 42,943,953 | 40,766,077 (94.9%) |
| IntS9 dsRNA Rep3 | 33,472,134 | 30,930,614 (92.4%) |

For each library, the raw reads were counted per gene based on Ensembl annotations (BDGP6.22.96) by htseq-count (version 0.11.2; parameters: -q -s reverse --nonunique all --additional-attr=gene_name -f bam) (Anders et al. 2015) and subjected to DESeq2 (version 1.24.0 under R version 3.6.1) (Love et al. 2014) to identify differentially expressed genes. For overlapping gene annotations with exactly the same coordinates, Ensembl gene ID and gene names were combined together. Genes with a fold change >1.5 and an adjusted *P* value <0.001 were considered to be significantly differentially expressed upon depletion of IntS9.

For the 409 up-regulated and 49 down-regulated genes identified by RNA-seq, Gene Ontology (GO) analysis was performed using the PANTHER14.1 statistical overrepresentation test (Mi et al. 2019). The annotation data set was GO biological process complete and the reference list was all genes in the *Drosophila melanogaster* database. The test type was Fisher's Exact and the false discovery rate was calculated for the correction.

### Supplemental Figure S1

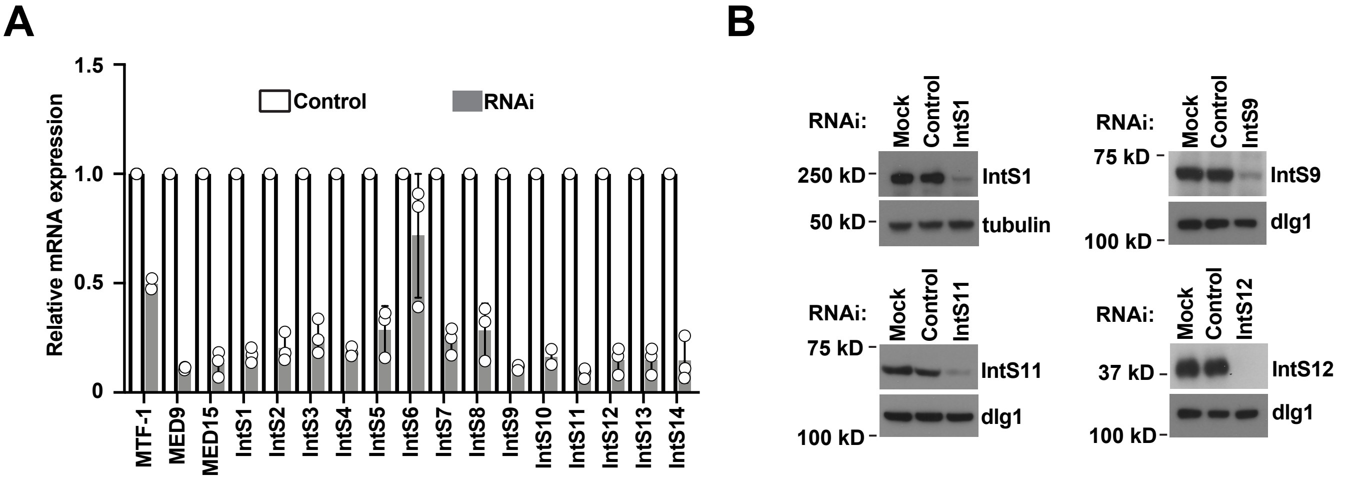

### Supplemental Figure S2

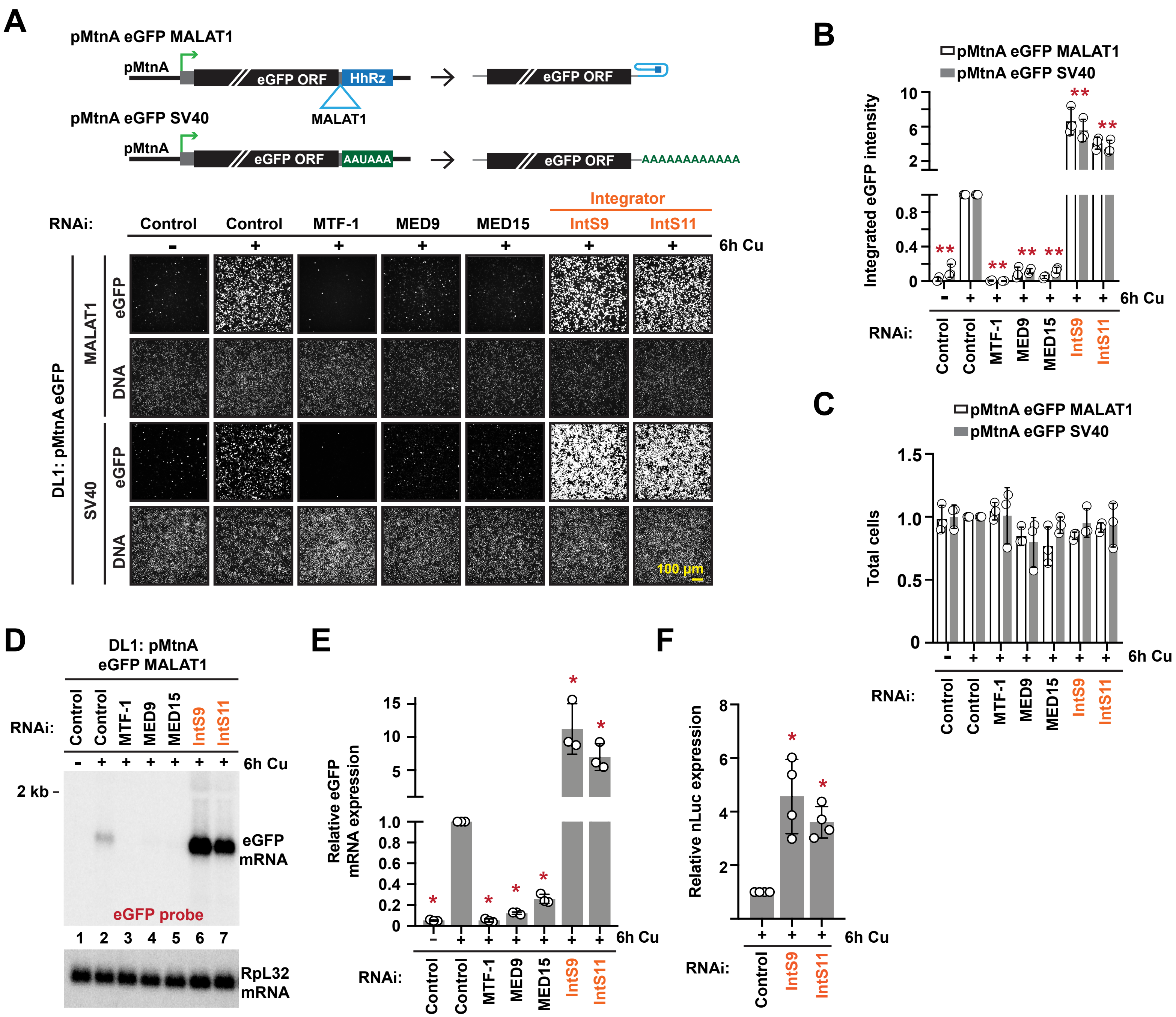

### Supplemental Figure S3

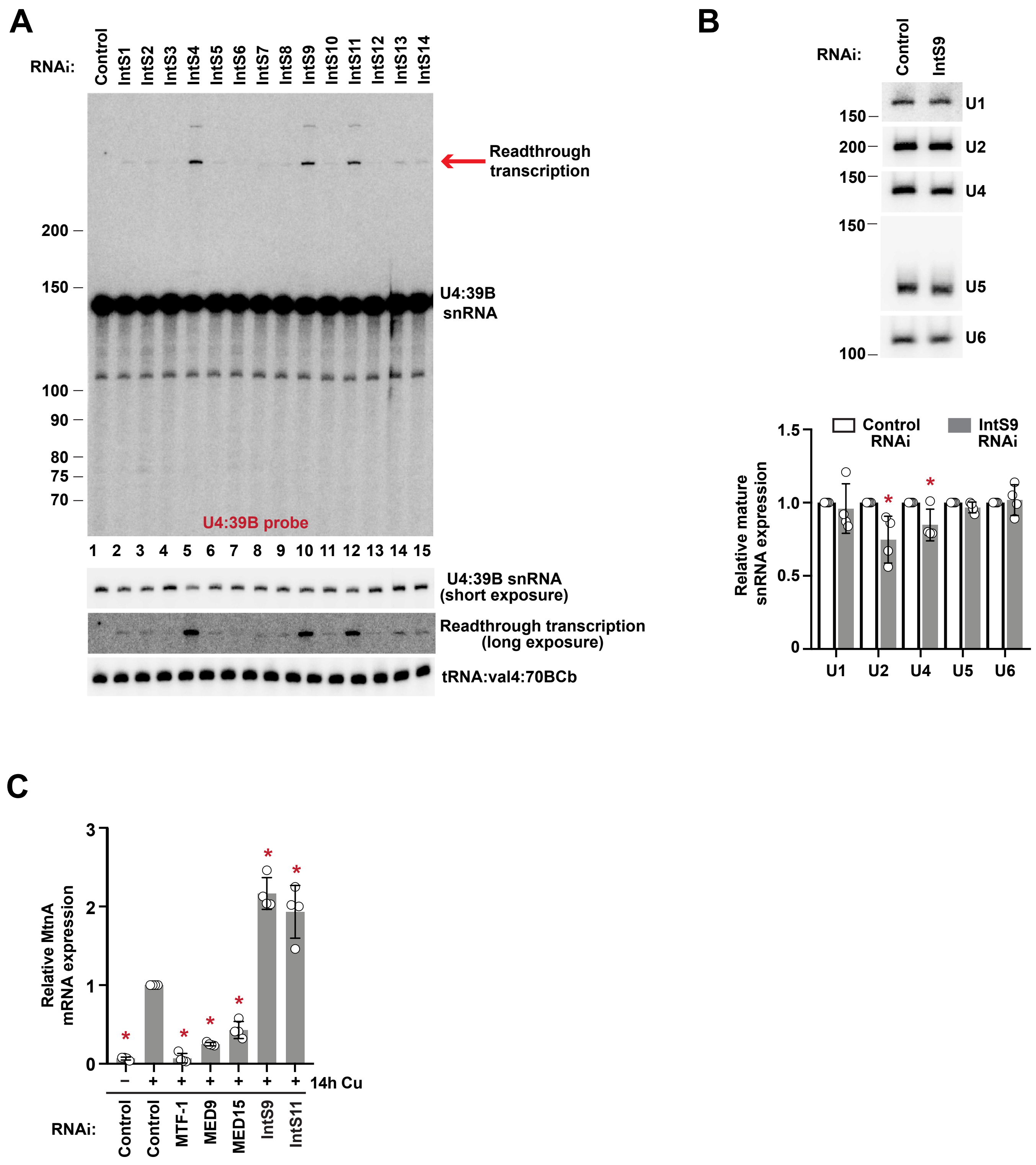

### Supplemental Figure S4

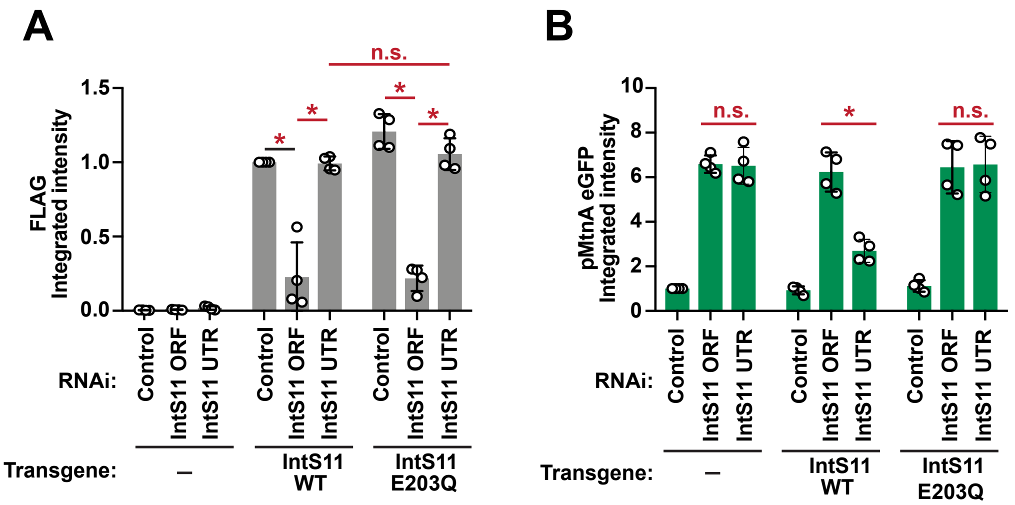

### Supplemental Figure S5

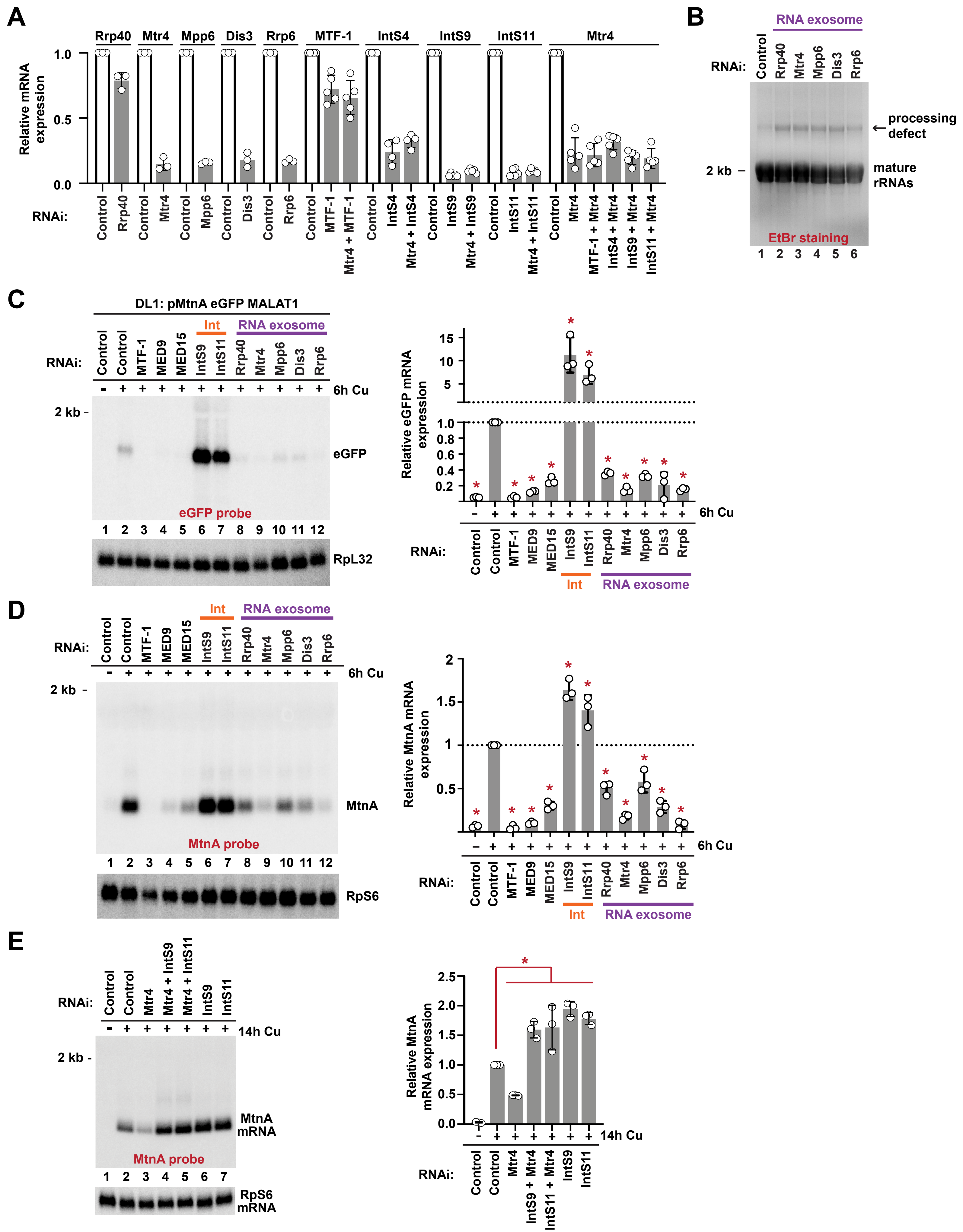

### Supplemental Figure S6

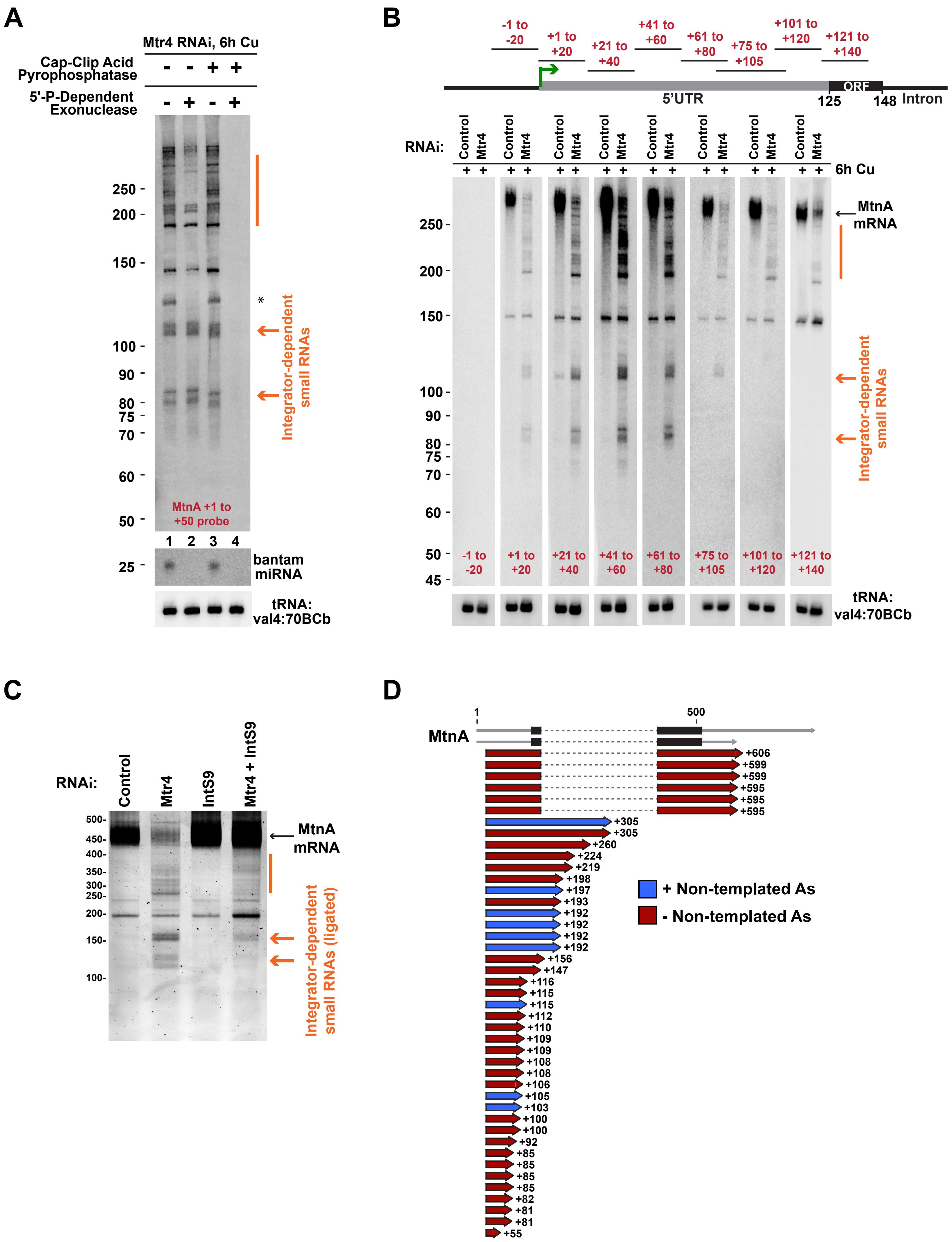

### Supplemental Figure S7

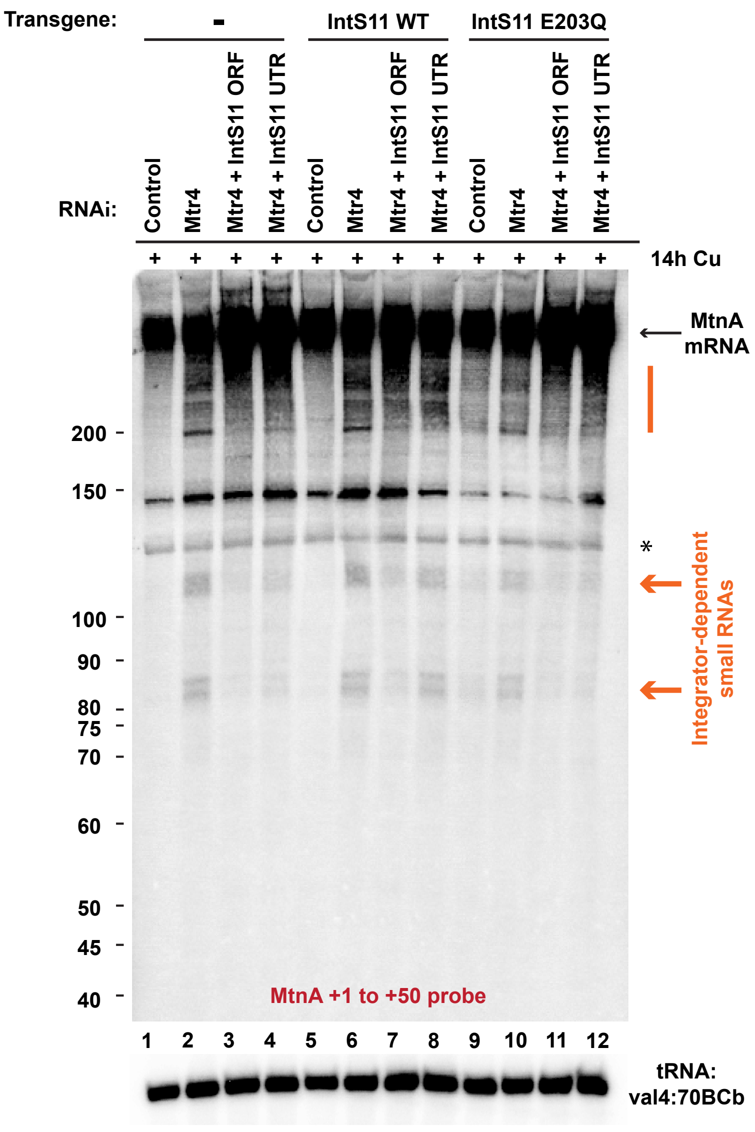

### Supplemental Figure S8

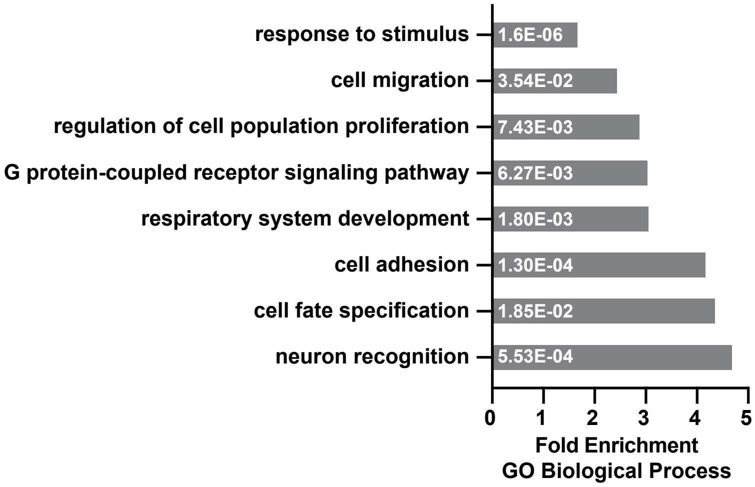

#### Supplemental Figure S9

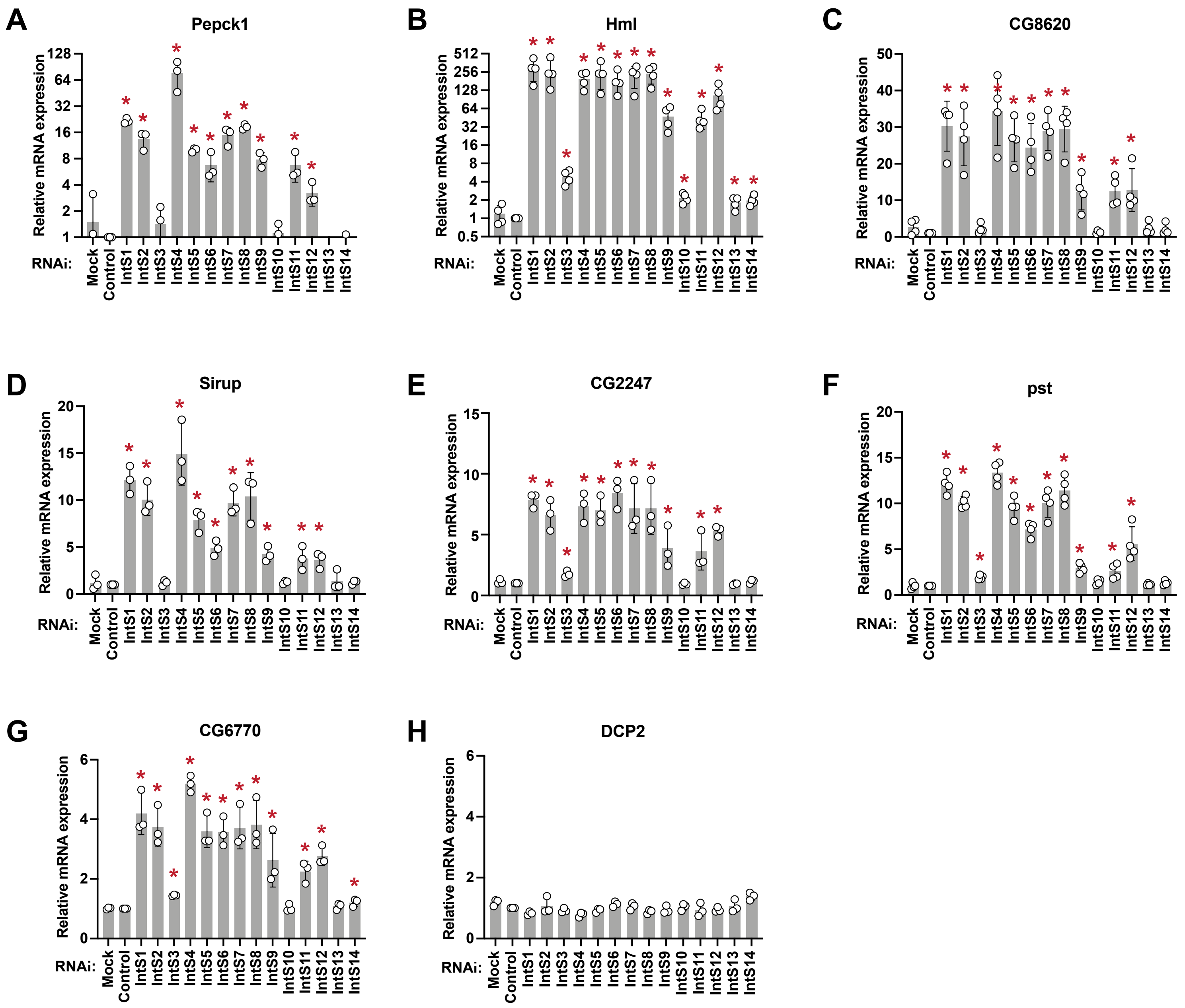

### Supplemental Figure S10

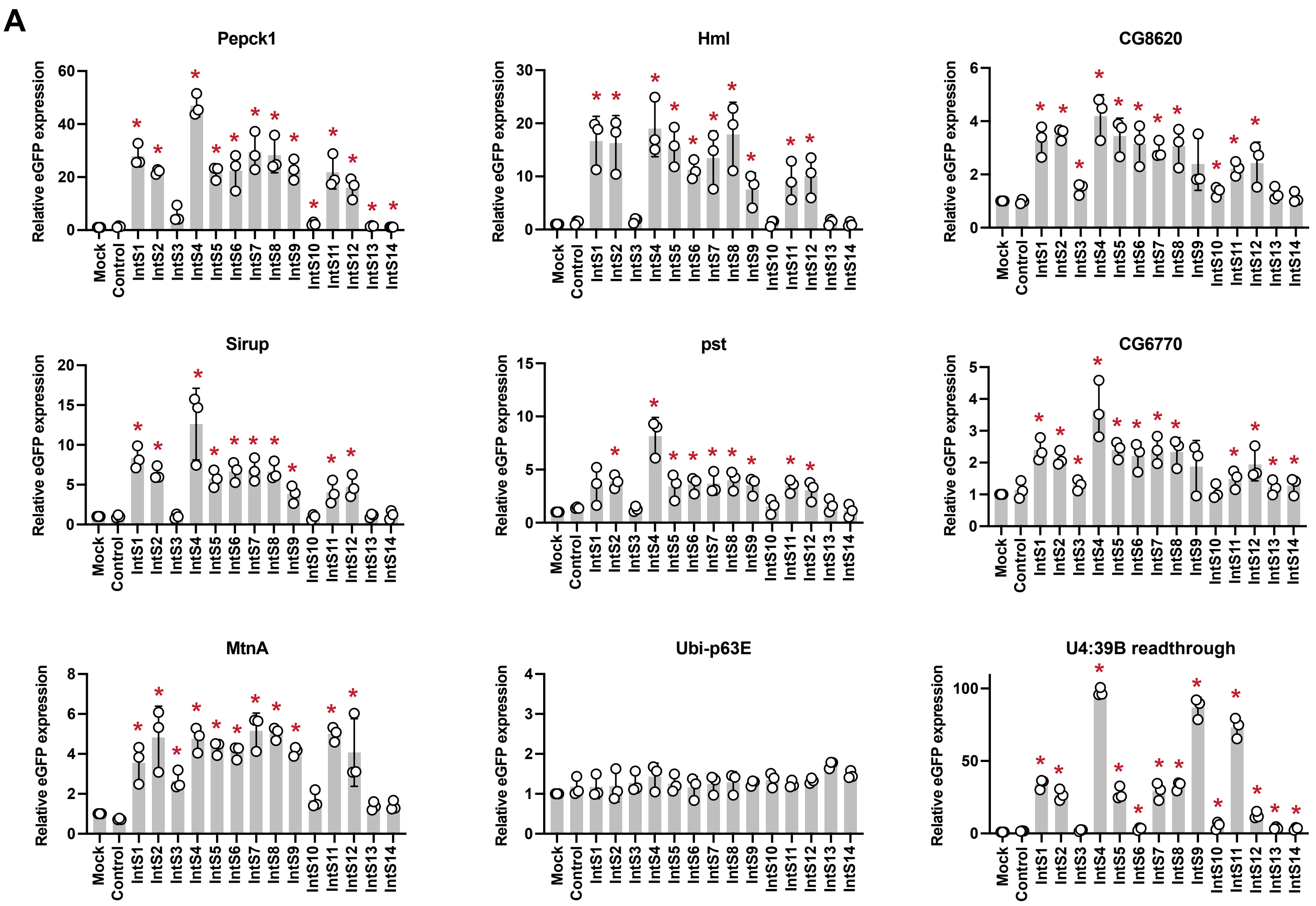


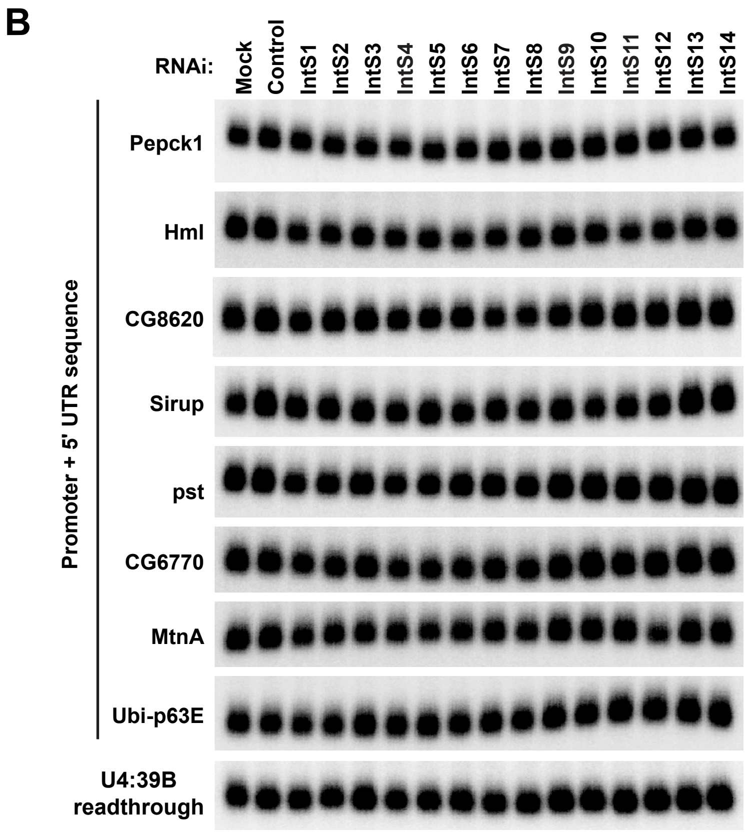

### Supplemental Figure S11

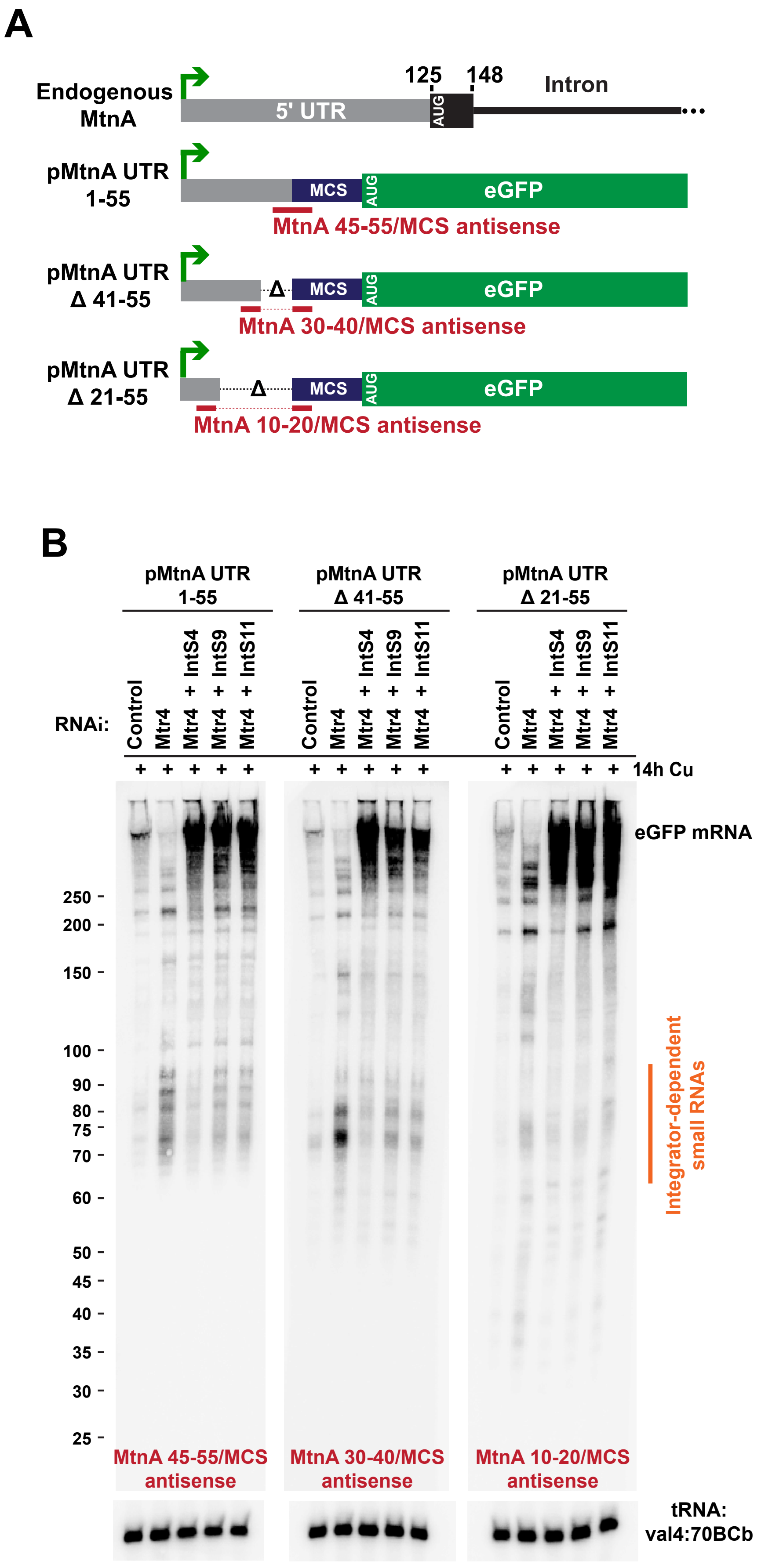

### Supplemental Figure S12

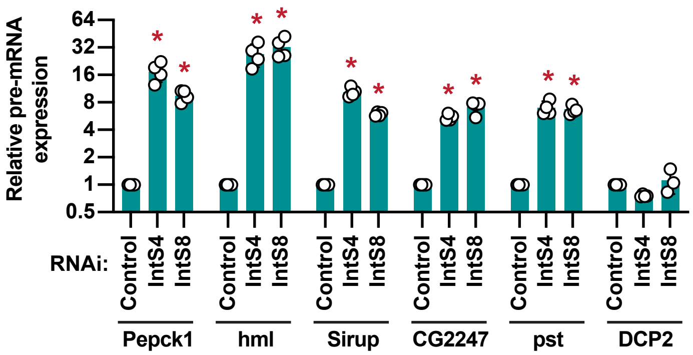

#### SUPPLEMENTAL FIGURE LEGENDS

**Supplemental Figure S1. Validation of Integrator depletion by RNAi in *Drosophila* DL1 cells.**

**(A)** RT-qPCR was used to quantify the efficiency of dsRNA-mediated depletion of target mRNAs. Data from 3 independent experiments were normalized to expression of DCP2 mRNA (for quantification of MTF-1, MED9, and MED15) or RpL32 mRNA (for quantification of all Integrator subunits) and are shown as mean ± SD. **(B)** DL1 cells were treated with dsRNAs for 3 days and then Western blotting was used to measure depletion of each Integrator protein subunit. dlg1 or α-tubulin were used as loading controls. Representative blots are shown.

**Supplemental Figure S2. Validation of the Integrator complex as an inhibitor of the MtnA promoter.**

**(A-C)** To determine if the mature eGFP mRNA must be processed at its 3' end in a particular way in order for it to be regulated by Integrator, DL1 cells stably maintaining an eGFP reporter ending in the MALAT1 triple helix or the SV40 polyadenylation signal were treated with dsRNAs for 3 days to induce RNAi and depletion of the indicated factors. $CuSO_4$ was added for the last 6 h. eGFP and DNA (Hoechst 33342) were then visualized **(A)**. Representative images are shown. The integrated eGFP intensity (amount of eGFP signal in each well divided by the number of cells) **(B)** and cell numbers **(C)** were quantified. Data are shown as mean ± SD, N=3. *$p<0.05$. Similar effects were observed regardless of the eGFP mRNA 3' end processing mechanism. (**D, E)** DL1 cells stably maintaining an eGFP reporter ending in the MALAT1 triple helix were treated with dsRNAs for 3 days to deplete the indicated factors and $CuSO_4$ was added for the last 6 h. Northern blots were used to examine eGFP mRNA expression **(D)** and

ImageQuant used to quantify signal from three independent blots **(E)**. Data are normalized to the copper-treated control (βgal dsRNA) samples and are shown as mean ± SD. * $p<0.05$. **(F)** To confirm that Integrator activity on the output of the MtnA promoter did not depend on the presence of a particular ORF, DL1 cells were first treated with the indicated dsRNA. After 24 h, cells were transfected with a nLuc reporter driven by the MtnA promoter and incubated for an additional 48 h. Cells were treated with $CuSO_4$ for the final 6 h and then nLuc activity was measured. Data are normalized to the copper-treated control (βgal dsRNA) samples and are shown as mean ± SD, N=4. *$p<0.05$.

**Supplemental Figure S3. Depletion of Integrator subunits results in readthrough transcription downstream of snRNAs and up-regulation of endogenous MtnA expression during copper stress.**

**(A)** DL1 cells were treated with dsRNAs for 3 days to induce RNAi and depletion of the indicated factors. $CuSO_4$ was added for the last 14 h. Northern blots were then used to examine expression of mature U4:39B snRNA as well as readthrough transcription downstream of this snRNA. A representative blot is shown. ImageQuant was used to quantify readthrough transcription (red arrow, quantified in **Fig. 1E** and **Fig. 5C**). **(B)** DL1 cells were treated for 3 days with a control (βgal) dsRNA or a dsRNA to deplete IntS9. Northern blotting was then used to measure expression of mature snRNAs. Representative blots are shown and ImageQuant was used to quantify transcript levels. Data are shown as mean ± SD, N=4. * $p<0.05$. Note: U6 snRNA biogenesis does not require the Integrator complex and serves as a loading control. **(C)** As in **Fig. 1F**, Northern blotting was used to measure expression of endogenous MtnA mRNA in

DL1 cells that had been treated with the indicated dsRNAs for 3 days and $CuSO_4$ for the last 14 h. Data are shown as mean ± SD, N=4. * $p<0.05$.

**Supplemental Figure S4. IntS11 endonuclease activity is required for Integrator to regulate the eGFP reporter driven by the MtnA promoter during copper stress.**

**(A, B)** DL1 cells were treated with the indicated dsRNAs. After 24 h, cells were transfected with plasmids expressing a FLAG-tagged IntS11 transgene and a plasmid expressing an eGFP reporter driven by the MtnA promoter. On the following day, cells were treated with $CuSO_4$ for 6 h and samples were processed. **(A)** Expression of the IntS11 transgenes was quantified by immunofluorescence using a FLAG antibody and automated microscopy. The integrated FLAG intensity (amount of FLAG signal in each image divided by the number of cells in each image) was measured and relative data are shown as mean ± SD, N=3. * $p<0.05$; n.s. = not significant. **(B)** Expression of eGFP protein was similarly quantified using automated microscopy. Data are shown as mean ± SD, N=4. * $p<0.05$; n.s. = not significant.

**Supplemental Figure S5. The RNA exosome acts on transcripts derived from the MtnA promoter.**

(**A)** DL1 cells were treated with dsRNAs (or combination of dsRNAs) for 3 days to induce RNAi and depletion of the indicated factor(s). RT-qPCR was then used to quantify the efficiency of dsRNA-mediated depletion of target mRNAs. Data from 3 or more independent experiments (as indicated) were normalized to DCP2 mRNA expression and are shown as mean ± SD. **(B)** Total DL1 RNA was resolved using a 1.2% formaldehyde agarose gel and ribosomal RNAs (rRNAs) were visualized by ethidium bromide (EtBr) staining. Depletion of RNA exosome components caused a defect in rRNA processing and accumulation of a higher molecular weight transcript (denoted by arrow), as expected. **(C)** DL1 cells stably maintaining an eGFP reporter ending in the MALAT1 triple helix were treated with the indicated dsRNAs for 3 days and $CuSO_4$ was added for the last 6 h. Northern blots were used to examine eGFP mRNA expression and ImageQuant used to quantify signal from independent blots. Data are normalized to the copper-treated control (βgal dsRNA) samples and are shown as mean ± SD, N=3. * $p<0.05$. **(D, E)** Representative Northern blots of endogenous MtnA mRNA in DL1 cells treated with $CuSO_4$ and the indicated dsRNAs. Data are normalized to the copper-treated control (βgal dsRNA) samples and are shown as mean ± SD, N=3. * $p<0.05$. Note that depletion of the exosome-associated RNA helicase Mtr4 significantly reduced expression from the MtnA promoter, but this effect was rescued upon co-depletion of Integrator subunits. The quantification of MtnA mRNA expression after depletion of RNA exosome components is also presented in **Fig. 3E**.

**Supplemental Figure S6. The MtnA small RNAs are capped, have the same TSS as full-length MtnA mRNA, and can be oligoadenylated.**

**(A)** DL1 cells were treated with a dsRNA to deplete Mtr4 for 3 days and $CuSO_4$ was added for the last 6 h. As indicated, 10 μg of isolated total RNA was then treated with a 5'-3'-dependent exonuclease (Terminator) and/or Cap-Clip Acid Pyrophosphatase, which removes 5' cap structures. Representative Northern blots are shown. Consistent with the MtnA small RNAs having a 5' cap, treatment with Cap-Clip Acid Pyrophosphatase caused the MtnA small RNAs to run slightly faster (Lane 3) and be susceptible to digestion by a 5'-3'-dependent exonuclease (Lane 4). The bantam microRNA was used as a positive control for exonuclease activity, whereas the highly structured tRNA lacks a 5' cap and is resistant to degradation. **(B)** Schematic of the MtnA locus and the oligonucleotide probes (top). DL1 cells were treated with dsRNAs for 3 days to induce RNAi and depletion of the indicated factors. $CuSO_4$ was added for the last 6 h. 10 µg of total RNA was probed by Northern blot analysis with the designated probes (bottom). Representative blots are shown. **(C, D)** DL1 cells were treated with dsRNAs for 3 days to deplete the indicated factors and $CuSO_4$ was added for the last 14 h. The 3' ends of endogenous MtnA transcripts were then cloned with a ligation-based 3' RACE approach. **(C)** Amplified products were visualized by ethidium bromide (EtBr) staining. **(D)** 3' RACE products from Mtr4 RNAi treated cells were cloned, subjected to Sanger sequencing, and aligned to the MtnA locus. For each clone, the length of the transcript is given (the annotated MtnA TSS is +1). A subset of the cloned transcripts was oligoadenylated at their 3' ends (blue).

**Supplemental Figure S7. The IntS11 endonuclease activity is required for expression of the MtnA small RNAs.**

Parental DL1 cells (Lanes 1-4) or DL1 cells stably expressing WT (Lanes 5-8) or catalytically inactive (Lanes 9-12) IntS11 transgenes were treated with the indicated dsRNAs for 3 days and $CuSO_4$ for the final 14 h. Transcripts generated from the MtnA locus were then analyzed by Northern blotting. Full-length MtnA mRNA (black arrow) and Integrator-dependent small RNAs (orange) are indicated. *, non-specific band. Representative blots are shown.

**Supplemental Figure S8. Gene Ontology (GO) analysis of Integrator regulated genes.**

The PANTHER overrepresentation test was performed on the 409 genes that were up-regulated upon IntS9 depletion. Fold enrichment is given for selected classes from the “GO Biological Process” annotation and represents the number of genes in the up-regulated gene set divided by the expected number of genes based on the entire *Drosophila* genome. The P value (noted in white) is adjusted to reflect the FDR. No significantly overrepresented classes were identified in the list of 49 genes that were down-regulated upon IntS9 depletion.

**Supplemental Figure S9. Up-regulation of mRNA expression upon depletion of Integrator subunits.**

DL1 cells were treated with the indicated dsRNAs. RT-qPCR was then used to quantify expression of the following mRNAs: **(A)** Pepck1, **(B)** Hml, **(C)** CG8620, **(D)** Sirup, **(E)** CG2247, **(F)** pst, **(G)** CG6770, and **(H)** DCP2 (which is not regulated by Integrator). Data are normalized to the control (βgal dsRNA) samples and are shown as mean ± SD, N≥3. * $p<0.05$. Note that the average mRNA expression value for each RNAi treatment is depicted in the heat map in **Fig. 5C**.

**Supplemental Figure S10. eGFP reporter genes driven by the exemplar promoters are regulated by Integrator.**

**(A)** As described in **Fig. 6**, DL1 cells were treated with the indicated dsRNAs and then transfected with the indicated plasmid. ImageQuant was used to quantify eGFP signal from three independent Northern blots. Data are normalized to the Mock treated samples and are shown as mean ± SD. * $p<0.05$. **(B)** Equal loading of RNA was verified for the Northern Blots shown in **Fig. 6** by re-probing the membranes for RpL32 mRNA.

**Supplemental Figure S11. Integrator cleavage of transcripts generated from the MtnA promoter is independent of the 5' UTR sequence.**

**(A)** DL1 cell lines that stably maintain eGFP reporters driven by the MtnA promoter with either 55 nt (pMtnA UTR 1-55), 40 nt (pMtnA UTR Δ41-55), or 20 nt (pMtnA UTR Δ21-55) of the MtnA 5' UTR were generated. Oligonucleotide probes for each construct are noted (red). **(B)** Each of the DL1 cell lines stably maintaining an eGFP reporter were then treated with the indicated dsRNAs for 3 days and $CuSO_4$ for the final 14 h. Northern blotting was then used to analyze small RNAs. Each of the eGFP reporters generated Integrator-dependent small RNAs that were stabilized upon Mtr4 depletion. Truncating the 5' UTR did not result in obvious shortening of the small RNAs, arguing against the presence of a 3′ box-like sequence element that dictates the Integrator cleavage sites.

**Supplemental Figure S12. Integrator regulates the transcription of protein-coding genes during cadmium stress.**

RT-qPCR was used to measure the pre-mRNA levels of the indicated transcripts after DL1 cells had been treated with dsRNAs for 3 days and $CdCl_2$ for the last 14 h. Data were normalized to RpL32 mRNA expression and are shown as mean ± SD, N≥3. * $p<0.05$.

#### SUPPLEMENTAL TABLE LEGENDS

**Supplemental Table S1. Negative and positive regulators of the pMtnA eGFP MALAT1 reporter.**

Gene hits from the RNAi screen are ordered by Robust Z score. "Negative regulators" are genes whose depletion resulted in increased expression of the reporter (Z score>1.3). "Positive regulators" are genes whose depletion resulted in decreased expression of the reporter (Z score<-1.3). Relative total cell numbers observed upon depleting each gene is also provided.

**Supplemental Table S2. Mapping of MtnA small RNA 3′ ends by RACE.**

DL1 cells were treated with a dsRNA to deplete Mtr4 for 3 days and $CuSO_4$ was added for the last 14 h. The 3' ends of endogenous MtnA transcripts were cloned and sequenced with a ligation-based 3' RACE approach. The MtnA genomic sequence (blue) is shown (exons in uppercase, introns in lowercase), starting at nt +22 from the TSS. The sequence and length of each 3′ RACE clone is given, with nucleotides that did not map to the *Drosophila* genome marked in red. A subset of 3′ RACE clones were spliced, with the exon-exon junction noted with a dash.

**Supplemental Table S3. *Drosophila* genes identified by RNA-seq as sensitive to depletion of IntS9.**

List of genes that are up-regulated (n=409) or down-regulated (n=49) after IntS9 RNAi as determined by RNA-seq. Threshold used to define IntS9-affected genes was fold change >1.5 and p<0.001. Replicate counts in DL1 cells treated with a control (βgal) dsRNA or a dsRNA to deplete IntS9 are shown for each transcript, along with the corresponding p-value used to identify differentially expressed genes.

**Supplemental Table S4. Oligonucleotide sequences.**

The oligonucleotide sequences used for Northern blots, RT-qPCR, ChIP-qPCR, 3' RACE, plasmid cloning, and dsRNA synthesis are provided.

**Supplemental Table S5. Antibodies for immunofluorescence, Western blots, and ChIP-qPCR.**

For each antibody used in this study, the source and experimental conditions are provided.
